# Cooperative thalamocortical circuit mechanism for sensory prediction errors

**DOI:** 10.1101/2023.07.12.548664

**Authors:** Shohei Furutachi, Alexis D. Franklin, Thomas D. Mrsic-Flogel, Sonja B. Hofer

## Abstract

The brain functions as a prediction machine, utilizing an internal model of the world to anticipate sensations and the outcomes of our actions. Discrepancies between expected and actual events, referred to as prediction errors, are leveraged to update the internal model and guide our attention towards unexpected events^1–10^. Despite the importance of prediction error signals for various neural computations across multiple brain regions, surprisingly little is known about the neural circuit mechanisms responsible for their implementation. Here we describe a thalamocortical disinhibitory circuit required for generating sensory prediction errors in mouse primary visual cortex (V1). Using calcium imaging with optogenetic manipulations as mice traverse a familiar virtual environment, we show that violation of animals’ predictions by an unexpected visual stimulus preferentially boosts responses of layer 2/3 V1 neurons most selective for that stimulus. Prediction errors specifically amplify the unexpected visual input, rather than representing a non-specific surprise or difference signal about how the visual input deviates from animals’ predictions. Selective amplification of unexpected visual input is implemented by a cooperative mechanism requiring thalamic input from the pulvinar, and cortical vasoactive-intestinal-peptide-expressing (VIP) inhibitory interneurons. In response to prediction errors, VIP neurons inhibit a specific subpopulation of somatostatin-expressing (SOM) inhibitory interneurons that gate excitatory pulvinar input to V1, resulting in specific pulvinar-driven response-amplification of the most stimulus-selective neurons in V1. Therefore, the brain prioritizes unpredicted sensory information by selectively increasing the salience of unpredicted sensory features through the synergistic interaction of thalamic input and neocortical disinhibitory circuits.

## Sensory prediction errors in visual cortex

Although our senses are continuously bombarded with inputs from the environment, only a subset of the sensory information is perceived or affects behavior. Our brains thus prioritize important sensory features among irrelevant ones^11^. Psychological and physiological studies indicate that the brain generates internal predictions about incoming sensory information and compares it with actual sensory inputs^5–10^, resulting in prediction errors when sensory inputs do not match internal predictions. Error signals could mediate prioritization of unexpected – and therefore possibly relevant – sensory inputs, and be used to update internal predictions^5–10^. Indeed, sensory prediction error signals have been observed in multiple cortical areas upon the violation of subjects’ predictions^9,10, 12–16.^ Despite their prevalence across the brain and importance for perception and learning, very little is known about what information is encoded by sensory prediction errors, how they affect cortical networks, and by which circuit mechanisms they arise.

To study the neural implementation of predictive processing in cortical sensory networks, we used a paradigm in which head-fixed, food-deprived mice running on a cylinder navigated a virtual corridor in which they developed spatial predictions about stimulus identity at particular locations along the corridor. The corridor walls displayed alternating grating patterns (grating A – grating B – grating A – grating B) separated by distinct landmarks (Fig. 1a). The gratings appeared abruptly when mice reached the corresponding position in the corridor and were presented at constant visual flow independent of mice’s running speed, to enable precise control over stimulus features and timing (see Methods). Upon reaching the reward zone at the end of the corridor, mice received a liquid food reward (see Methods) and their position was reset to the beginning of the corridor, starting a new trial. Mice traversed the corridor many times for five days of training (90 ± 48 trials (traversals) per day, 59 ± 21 seconds per trial; mean ± standard deviation (SD)) during which the sequence of the gratings was identical on every trial. On day six (C session), the identity of the stimulus at the fourth position changed in a subset of trials: a novel grating stimulus C was first shown instead of the second grating B in 10% of trials (block 1, 160 trials in total, Fig. 1a). Subsequently, grating C was shown at the fourth location in all trials (block 2, 40 trials). Previous studies using similar paradigms showed that mice form predictions of which stimuli to expect at specific locations in the corridor^14, 17^. Accordingly, we found that mice interrupted their running behavior when their expectations were violated by encountering grating C (Extended Data Fig. 1), similar to the interruption of ongoing actions by sensory surprise observed in humans^18^.

**Fig. 1.**
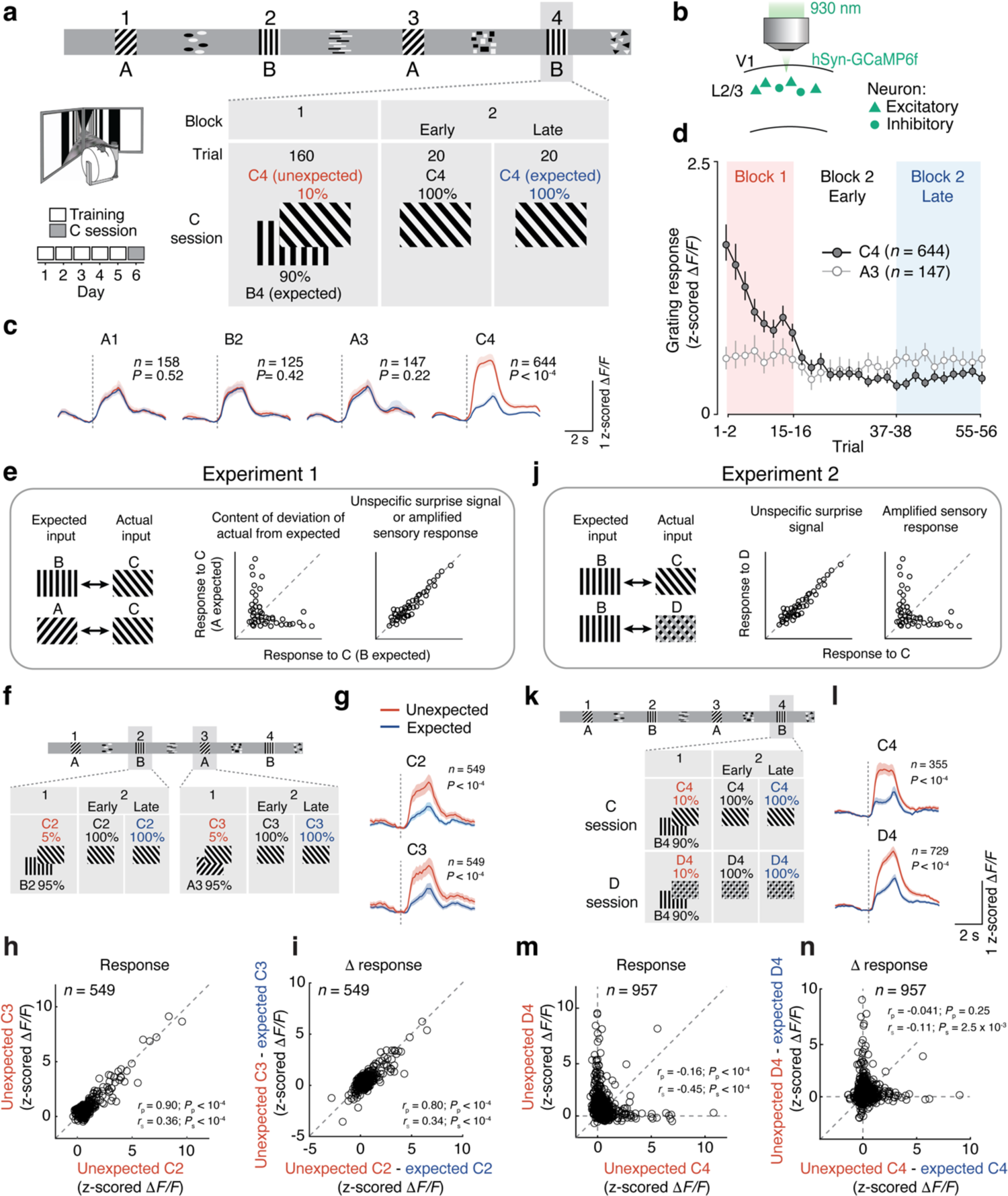
Prediction errors amplify unexpected visual information. **a**, Schematic of the structure of the virtual corridor. After five training sessions with grating B at location 4 (corridor as shown on top), grating C was shown in 10% of trials at position 4 in block 1 (referred to as unexpected C4) and 100% of trials in block 2 (referred to as expected C4 in the second half of block 2). **b**, Schematic of the two-photon calcium imaging approach. The activity of V1 layer 2/3 cells was recorded. **c**, Average calcium responses to gratings A1, B2, A3 and C4 in unexpected C trials (red, traversals with grating sequence A1 – B2 – A3 – C4 in block 1) and in expected C trials (blue, traversals in late block 2). V1 neurons responsive to the presented grating stimulus in unexpected C trials, expected C trials or both were included (see Methods). Dotted vertical lines depict grating onsets. Lines and shaded regions are mean and bootstrap 95% confidence interval (CI), respectively. A1: *n* = 158, *P* = 0.52; B2: *n* = 125, *P* = 0.42; A3: *n* = 147, *P* = 0.22; C4: *n* = 644, *P* < 1.0 × 10^-4^; A1, B2, A3, C4 responsive cells from 9 mice; Wilcoxon signed-rank test. See also Extended Data Fig. 2 for combined responses of all grating-responsive neurons. **d**, Average calcium responses to C4 (dark gray) and A3 (light gray) during C trials across trials and blocks. Symbols and error bars depict mean and bootstrap 95% CI. **e**, Thought experiment to disambiguate different hypotheses of what information prediction errors represent. **f**, Experimental design. Grating C was presented at position 2 (C2) or at position 3 (C3) in 5% of trials each in block 1. **g,** Average calcium responses to unexpected (red) and expected (blue) C2 (top, *n* = 549 from 9 mice, *P* < 1.0 × 10^−4^, Wilcoxon signed-rank test) and C3 (bottom, *n* = 549 from 9 mice, *P* < 1.0 × 10^−4^). Cells responsive to unexpected or expected C2 or C3 were pooled. Lines and shading indicate mean and bootstrap 95% CI. **h**, Responses to unexpected grating stimulus C2 plotted against responses to unexpected C3 for individual V1 layer 2/3 neurons (*r*_p_ = 0.90, *r*_s_ = 0.36; P_p_ < 1.0 × 10^−4^, P_s_ < 1.0 × 10^−4^; Pearson and Spearman correlation, respectively; *n* = 549 cells from 9 mice). **i**, Difference in response strength between grating responses to unexpected and expected C2 (prediction error signal) plotted against response strength difference between unexpected and expected C3 responses for individual V1 layer 2/3 neurons (*r*_p_ = 0.80, *r*_s_ = 0.34; *P*_p_ < 1.0 × 10^−4^, *P*_s_ < 1.0 × 10^−4^; Pearson and Spearman correlation, respectively). **j**, Similar to e, but for a second thought experiment. **k**, Experimental design. Gratings C or D were presented at position 4 (C4 and D4) in 10% of trials in different sessions (C and D sessions, respectively). **l**, Average V1 calcium responses to unexpected (red) and expected (blue) grating C4 (top, *n* = 355 from 5 mice, *P* < 1.0 × 10^−4^) and D4 (bottom, *n* = 729 from 5 mice, *P* < 1.0 × 10^−4^). Wilcoxon signed-rank test. Lines and shading indicate mean and bootstrap 95% CI. **m**, Responses to unexpected stimulus C4 plotted against responses to unexpected D4 for individual V1 layer 2/3 neurons (*r*_p_ = −0.16, *r*_s_ = − 0.45; *P*_p_ < 1.0 × 10^−4^, *P*_s_ < 1.0 × 10^−4^; Pearson and Spearman correlation, respectively; *n* = 957 cells from 5 mice). **n**, Difference in response between grating responses to unexpected C4 and expected C4 stimulus plotted against response difference between unexpected D4 and expected D4 stimulus for individual V1 layer 2/3 neurons (*r*_p_ = −0.041, *r*_s_ = −0.11; *P*_p_ = 0.25, *P*_s_ = 2.5 × 10^−3^; Pearson and Spearman correlation, respectively). See also Extended Data Fig. 1 and 2.

We recorded neural activity of layer 2/3 neurons in V1 using two-photon calcium imaging (Fig. 1b, see Methods)^19^, and observed a stronger response to a visual stimulus that was novel and therefore unexpected (grating C in block 1) compared to the same stimulus when it was expected (grating C in second half of block 2, P < 1.0 × 10^−4^, Wilcoxon signed-rank test, Fig. 1c, Extended Data Fig. 2a, b), consistent with previous studies in humans, non-human primates and rodents^9,10,12–14, 16, 20–24.^ This difference in neural responses could not be explained by a drift in general behavioral state, such as arousal or task engagement across the imaging session, as responses to expected grating stimuli A and B were constant throughout the session (Fig. 1c, d, all *P* > 0.05, see also Extended Data Fig. 2a, b). The increased response to unexpected visual stimuli could also not be accounted for by changes in the animal’s motor behavior or a visuomotor mismatch signal caused by the mouse’s deceleration (Extended Data Fig. 1). Specifically, there was no difference in mice’s pupil size or licking behavior, and grating C response strength was not correlated with mice’s running speed or stimulus-induced deceleration (Extended Data Fig. 1).

**Fig. 2.**
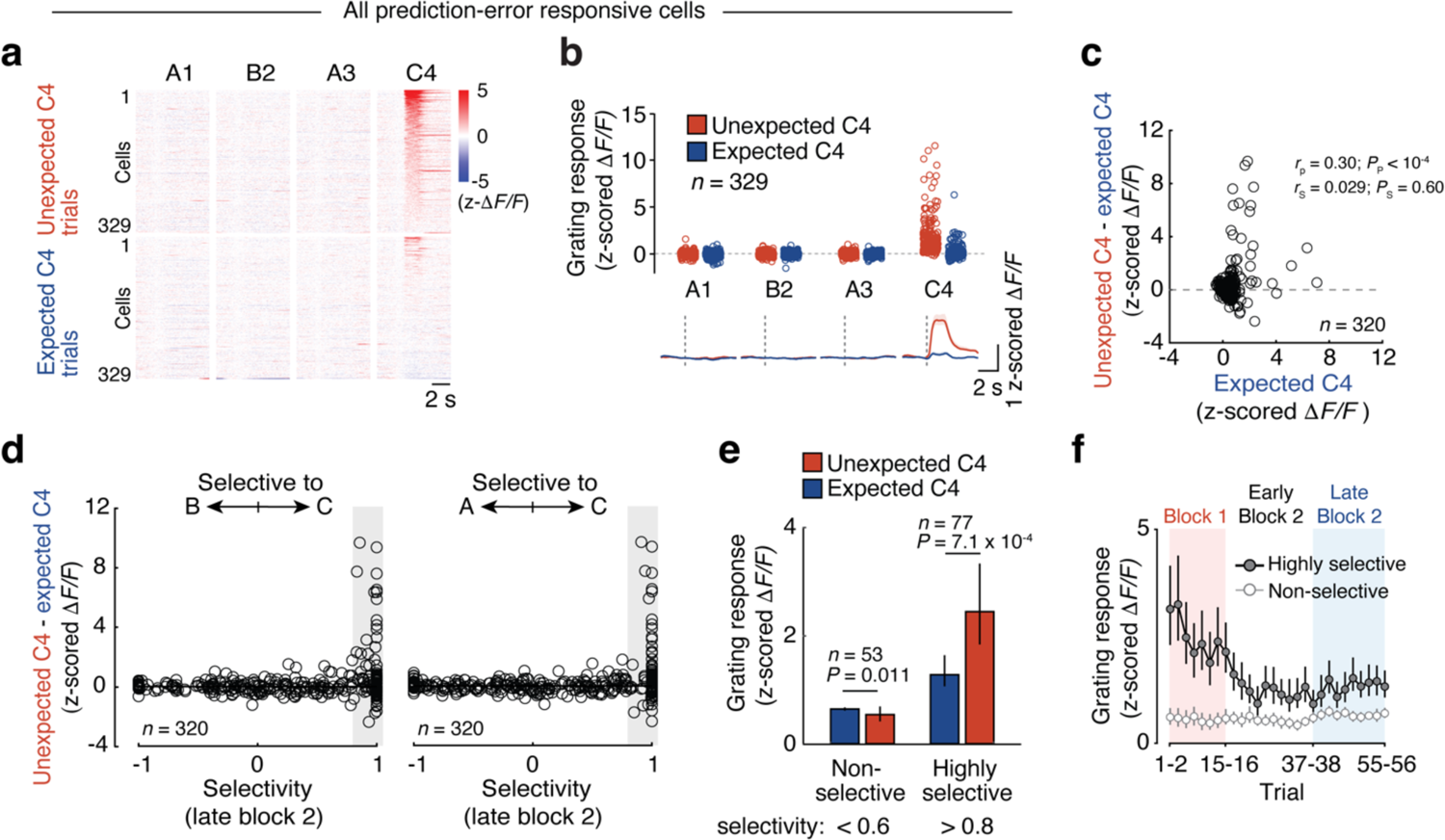
Prediction error specifically boosts neurons most selective to the unexpected visual stimulus. **a**, Trial-averaged responses of all prediction-error responsive V1 neurons (individual rows, *n* = 329 cells, 9 mice) to gratings A1, B2, A3 and C4 in unexpected C4 trials (top, block 1) and expected C4 trials (bottom, late block 2), sorted by response to unexpected C4. **b**, Top: Average response strength for individual prediction-error responsive cells (individual dots, *n* = 329 cells, 9 mice) to gratings A1, B2, A3 and C4 in unexpected C4 (red) and expected C4 (blue) trials. Bottom: mean calcium responses of all neurons above, shading indicates bootstrap 95% CI. **c**, Difference in response strength between unexpected (block 1) and expected C4 (late block 2) for all grating-responsive cells in late block 2, plotted against response to expected C4 in late block 2 for individual neurons (*r*_p_ = 0.30, *r*_s_ = 0.029; *P*_p_ < 1.0 × 10^−4^, *P*_s_ = 0.60; Pearson and Spearman correlation, respectively; *n* = 320, 9 mice). **d**, Left: difference in response strength between unexpected and expected C4 responses of individual neurons, plotted against their response selectivity to stimulus C vs. stimulus B in late block 2 (the difference in response strength between expected C4 and B2 divided by the sum of responses to both stimuli) for all neurons responsive to at least one of the grating stimuli in late block 2. −1 indicates only responsive to B, +1 only responsive to C, and 0 equal responses to both. Right: same as on the left but for response selectivity to stimulus C vs. stimulus A in late block 2. **e**, Mean responses to expected (blue) and unexpected (red) grating C4, of non-selective (selectivity towards C, compared to B < 0.6, left, *n* = 53; *P* = 0.011; Wilcoxon signed-rank test) and highly selective (selectivity towards C, compared to B > 0.8, right, *n* = 77; *P* =7.1 × 10^−4^) grating C4 responsive cells in late block 2. Error bars are 95% bootstrap CI. **f**, Mean calcium responses to stimulus C4 over all trials in the imaging session of highly selective (dark gray, *n* = 77) and non-selective (light gray, *n* = 53) grating C4 responsive cells in late block 2. Responses were averaged over two trials. Error bars are bootstrap 95% CI. See also Extended Data Fig. 3 and 4.

Neural responses to grating C strongly decreased over time as mice encountered the visual stimulus more often and asymptoted within several trials in block 2 when grating C was encountered in every trial (Fig. 1d). This gradual decrease in response cannot simply be explained by visual adaptation to repetitive stimuli, as grating C was only presented every 448 ± 364 (mean ± SD) seconds in block 1, due to the considerable length of the virtual corridor. Importantly, responses also significantly increased when a known visual stimulus A was presented at an unexpected location in the corridor (Extended Data Fig. 3; *P* = 0.0020 for highly selective cells), or when an expected stimulus was omitted (Extended Data Fig. 2e, f; *P* < 1.0 × 10^−4^ for grating omission)^14^. The elevated neural response to an unexpected stimulus is therefore most consistent with a prediction error signal. Indeed, the gradual decrease and eventual cessation of the prediction error signal after repeated exposure to the novel stimulus at the same location indicates that animals learn to update their spatial expectations about stimulus identity over time.

**Fig. 3.**
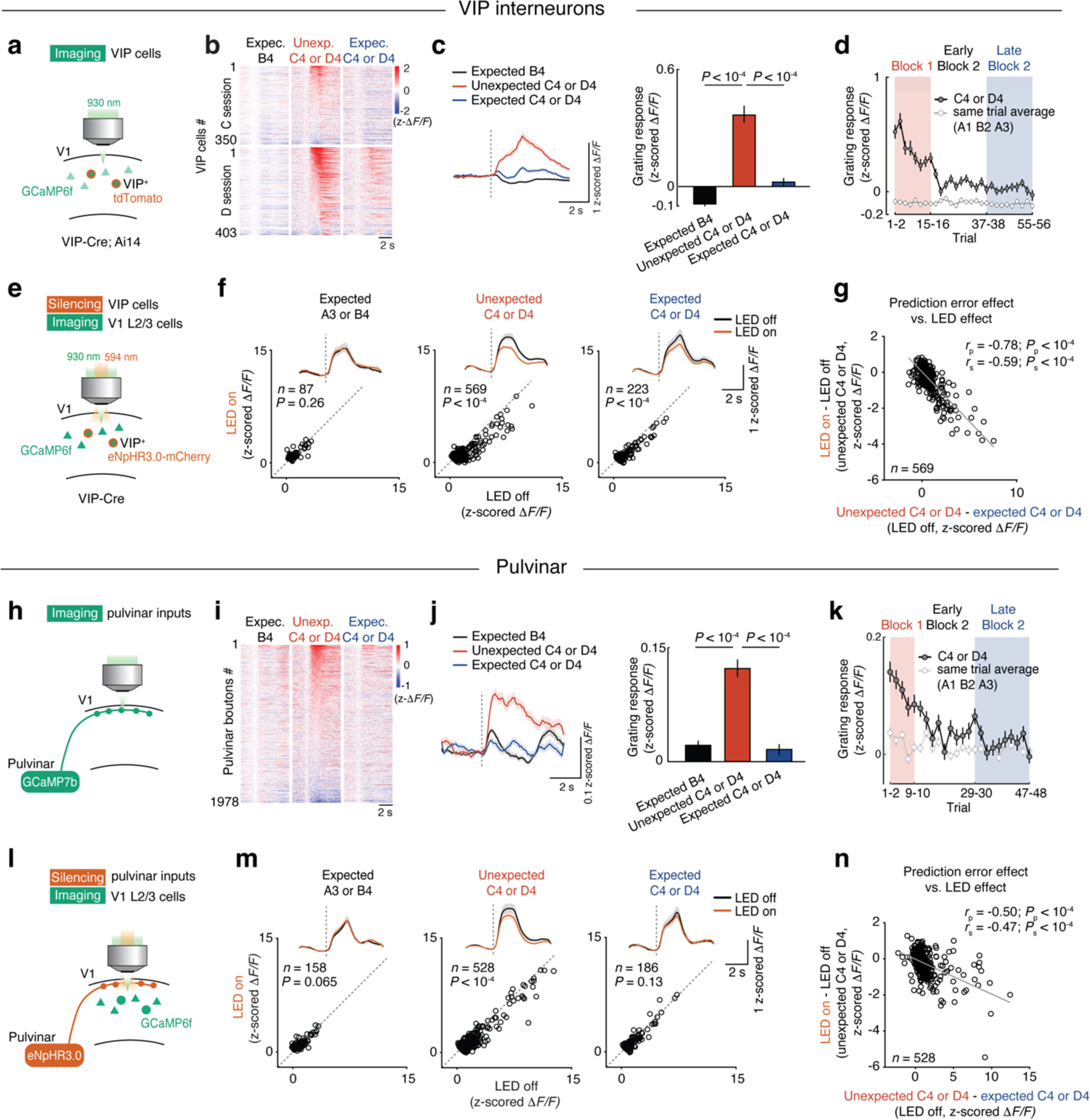
Activity of VIP interneurons and pulvinar input is required for prediction error signals in V1. **a,** Schematic of the experimental design. Calcium activity of VIP cells in V1 layer 2/3 was recorded during the experiment depicted in Fig. 1k. **b**, Single-cell responses for all VIP cells (individual rows) in the C session (top, *n* = 350 cells from 7 mice) and D session (bottom, *n* = 403 cells from 7 mice) to expected B4 (left), unexpected C4 or D4 (middle, block 1) and expected C4 or D4 (right, late block 2), sorted by response strength to unexpected C4 or D4. 290 cells were recorded in both sessions. **c**, Cell- and trial-averaged calcium responses of all VIP cells to expected B4 (black), unexpected C4 or D4 (red) and expected C4 or D4 (blue). All VIP cells from C session (*n* = 350) and D session (*n* = 403) were pooled. Lines and bars are mean, shading and error bars indicate bootstrap 95% CI. Responses to B4 vs C4/D4 in block 1: *P* < 1.0 × 10^−4^; C4/D4 in block 1 vs C4/D4 in late block 2: *P* < 1.0 × 10^−4^, Wilcoxon signed-rank test with Bonferroni correction. **d,** Average calcium responses of all VIP cells to grating C4 or D4 (dark gray) and other gratings in the same trial (average of A1, B2 and A3, light gray) over time. Symbols and error bars are mean and bootstrap 95% CI. **e**, Schematic of the experiment. Calcium activity of V1 layer 2/3 cells was recorded while VIP cells were optogenetically silenced during the experiment depicted in Fig. 1k (see Methods). VIP cell silencing started at the onset of grating stimuli and lasted for 3 s. **f**, Top: cell- and trial-averaged responses of neurons significantly responsive to the presented grating stimuli to expected grating A3 or B4 (left, *P* = 0.26, Wilcoxon signed-rank test, *n* = 87 cells, 7 mice), unexpected grating C4 or D4 (middle, *P* < 1.0 × 10^−4^, *n* = 569) and expected grating C4 or D4 (right, *P* < 1.0 × 10^−4^, *n* = 223) with (amber) or without (black) VIP silencing. Lines and shading are mean and bootstrap 95% CI. Bottom: responses of individual neurons to the grating stimulus indicated above during VIP silencing (LED on), plotted against responses to the same stimulus in control trials (LED off). **g**, Effect of VIP neuron silencing (LED on - LED off during unexpected grating C4 or D4) plotted against the strength of prediction error signals (response to unexpected C4 or D4 - response to expected C4 or D4). Pearson correlation: *r*_p_ = −0.78, *P*_p_ < 1.0 × 10^−4^; Spearman correlation: *r*_s_ = −0.59, *P*_s_ < 1.0 × 10^−4^. **h-k**, Same as a-d, but for calcium responses of pulvinar axonal boutons in V1 (see Methods). **h**, Calcium activity of axonal boutons of pulvinar projections was recorded in V1 layer 1. **i**, Single-bouton responses for all pulvinar axonal boutons (individual rows) in the C or D session (*n* = 1,978 boutons, 10 sessions from 7 mice) to expected B4 (left), unexpected C4 or D4 (middle, block 1) and expected C4 or D4 (right, late block 2), sorted by response strength to unexpected C4 or D4. **j**, Bouton- and trial-averaged calcium responses of all pulvinar boutons to expected grating B4 (black), unexpected grating C4 or D4 (red) and expected grating C4 or D4 (blue). Lines and bars are mean, shading and error bars indicate bootstrap 95% CI. Responses to B4 vs C4/D4 in block 1: P < 1.0 × 10^−4^; C4/D4 in block 1 vs C4/D4 in late block 2: P < 1.0 × 10^−4^, Wilcoxon signed-rank test with Bonferroni correction. **k,** Average calcium responses of all pulvinar boutons to grating C4 or D4 (dark gray) and other gratings in the same trial (average of A1, B2 and A3, light gray) over time. Symbols and error bars are mean and bootstrap 95% CI. **l-n**, Same as e-g, but with optogenetic silencing of pulvinar axons. **l**, The activity of V1 layer 2/3 cells was recorded while pulvinar axons in V1 were optogenetically silenced (see Methods). **m**, Top: cell- and trial-averaged responses of neurons significantly responsive to the presented grating stimuli to expected grating A3 or B4 (left, *P* = 0.065, Wilcoxon signed-rank test, *n* = 158 cells, 9 sessions from 8 mice), unexpected grating C4 or D4 (middle, *P* < 1.0 × 10^−4^, *n* = 528) and expected grating C4 or D4 (right, *P* = 0.13, *n* = 186) with (amber) or without (black) silencing of pulvinar axons. Lines and shading are mean and bootstrap 95% CI. Bottom: responses of individual neurons to the grating stimulus indicated above during silencing of pulvinar axons (LED on), plotted against responses to the same stimulus in control trials (LED off). **n**, Effect of silencing of pulvinar axons (LED on - LED off during unexpected grating C4 or D4) plotted against the strength of prediction error signals (response to unexpected C4 or D4 - response to expected C4 or D4). Pearson correlation: *r*_p_ = −0.50, *P*_p_ < 1.0 × 10^−4^; Spearman correlation: *r*_s_ = −0.47, *P*_s_ < 1.0 × 10^−4^. See also Extended Data Fig. 5, 6 and 7.

### Nature of sensory prediction error signals

What information sensory prediction errors represent is currently unclear. According to the theory of predictive coding, prediction errors have been proposed to encode the difference between predicted and actual visual input (encoding the content of how the actual visual input is different from predictions)^5–8^. However, error responses could also represent a more unspecific surprise signal (encoding only the magnitude of the deviation), or could enhance the representation of unpredicted sensory input (encoding the content of the actual input). We designed further experiments to disambiguate between these options. First, in a small subset of trials, we presented grating C at one of two different locations in the corridor, at which either grating B (position 2) or grating A (position 3) were expected (Experiment 1, Fig. 1e, f). Grating C elicited a stronger response in V1 in either location when it was unexpected (Fig. 1g). In these two instances the actual visual stimulus is the same, but animals’ predictions are likely to be different. If the prediction error signal represents the predicted stimulus and/or how the actual stimulus deviates from this prediction, V1 responses should differ to grating C at the two different locations. However, the responses of V1 neurons to the unexpected grating C in the two locations were notably similar (Fig. 1h, i; *r*_p_ = 0.90, *r*_p_ = 0.80; *P*_p_ < 1.0 × 10^−4^, *P*_p_ < 1.0 × 10^−4^; *r*_s_ = 0.36, *r*_s_ = 0.34; *P*_s_ < 1.0 × 10^−4^, *P*_s_ < 1.0 × 10^−4^; for h and i, respectively), indicating that – at least at the level of individual neurons in V1 – the sensory prediction error signal contains little information about how the actual input differs from predictions.

Next, we tested if the prediction error signal represents the actual visual input or instead a non-specific surprise or motor-related signal (Experiment 2, Fig. 1j, k). To this end we introduced an additional unexpected visual stimulus D that was presented at corridor position 4 in a subset of trials in a separate imaging session of the same neuronal populations (Fig. 1j, k). Both stimuli C and D evoked strong prediction error responses when they were unexpected (Fig. 1l, Extended Data Figs. 2c, d and 4). Neural responses to C and D should be similar if they simply represented a non-specific surprise signal, or activity related to surprise-triggered movement, such as deceleration in response to an unexpected stimulus. However, most neurons responded strongly to only one of the two unexpected stimuli, V1 population responses to these stimuli were thus different and specific to stimulus features (Fig. 1m, n).

**Fig. 4.**
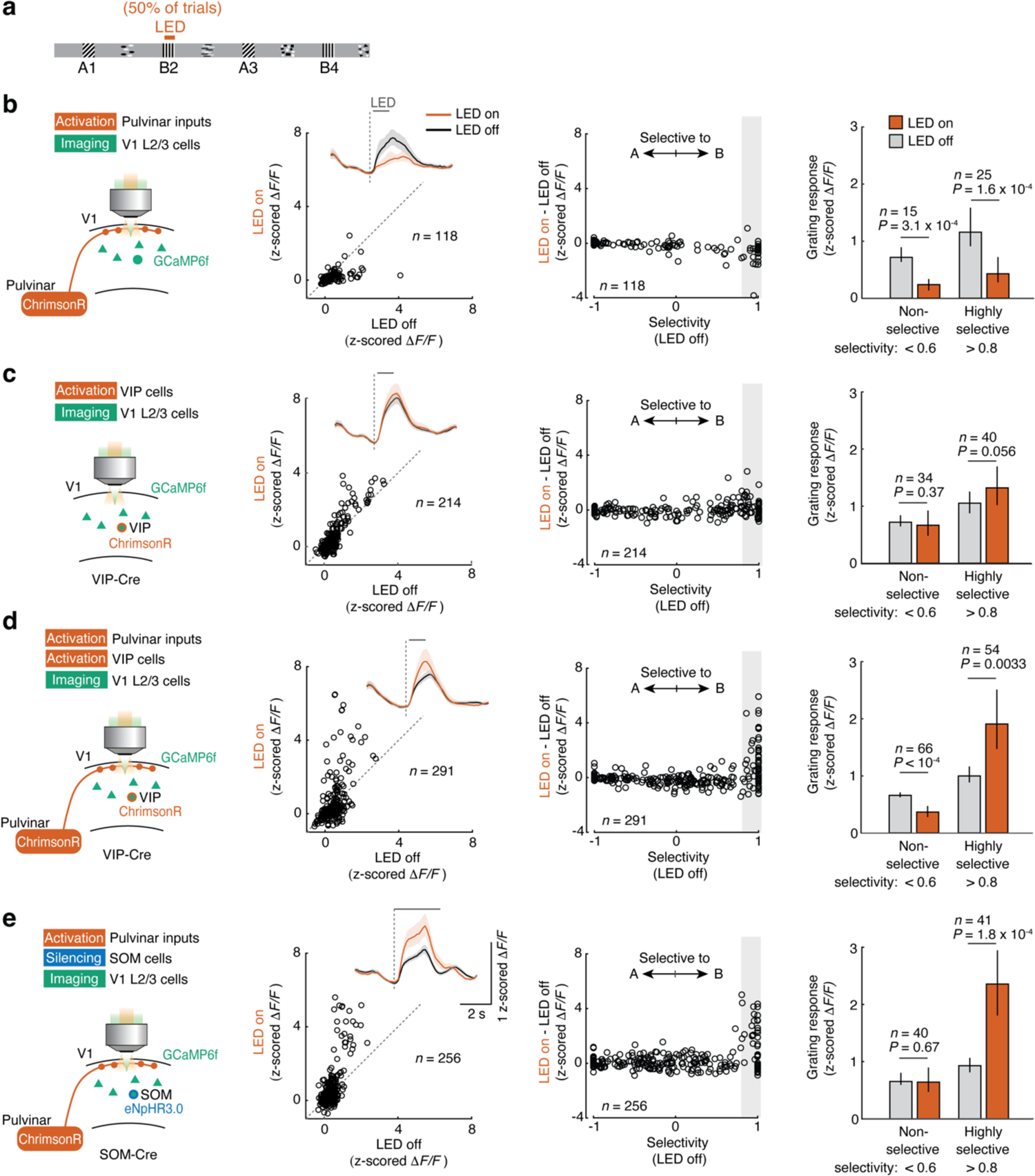
Neocortical disinhibition and pulvinar inputs act synergistically. **a**, Schematic of the experimental design. After training in the virtual corridor (stimuli A-B-A-B), optogenetic manipulation was paired with grating B2 in 50% of trials. **b**, Left: schematic of the experimental design. The activity of V1 layer 2/3 cells was recorded while pulvinar axons were optogenetically stimulated. Stimulation started 0.1 s after grating onset and lasted for 1 s (see methods). Second column: responses of individual V1 neurons with and without pulvinar axonal stimulation (LED on vs LED off). *n* = 118 grating A or B responsive cells, 6 mice, *P* < 1.0 × 10^−4^, Wilcoxon signed-rank test. Inset: cell-averaged calcium responses with (amber) or without (black) optogenetic stimulation. Lines and shaded regions are mean and bootstrap 95% CI, respectively. Third column: effect of optogenetic stimulation (difference of response to grating B2 with and without laser stimulation) plotted against response selectivity (difference in response strength stimulus B and A divided by the sum of responses) of individual V1 neurons. Right: calcium response strength to grating stimulus B2 of non-selective (selectivity B vs A < 0.6, left, *n* = 15, *P* = 3.1 × 10^-4^) and highly selective (selectivity B vs A > 0.8, right, *n* = 25, *P* = 1.6 × 10^-4^) grating B2 responsive cells in V1 layer 2/3 with (amber) or without (gray) optogenetic stimulation. Bars and error bars are mean and bootstrap 95% CI. **c**, Same as b, but with optogenetic stimulation of VIP cells. Left: the activity of V1 layer 2/3 cells was recorded while VIP cells were optogenetically stimulated for 1 s starting 0.1 s after grating onset. Second column: *n* = 214 cells from 6 mice, *P* = 0.43, Wilcoxon signed-rank test. Third column: *n* = 214 cells from 6 mice. Right: *n* = 34 and 40; *P* = 0.37 and *P* = 0.056; non-selective and highly selective cells; Wilcoxon signed-rank test. **d**, Same as b, but with simultaneous optogenetic stimulation of pulvinar axons and VIP cells. Left: the activity of V1 layer 2/3 cells was recorded while pulvinar axons and VIP cells were optogenetically co-stimulated for 1 s starting 0.1 s after grating onset. Second column: *n* = 291 cells from 8 mice, *P* = 0.037, Wilcoxon signed-rank test. Third column: *n* = 291 cells from 8 mice. Right: *n* = 66 and 54; *P* < 1.0 × 10^-4^ and *P* = 3.3 × 10^-3^; non-selective and highly selective cells; Wilcoxon signed-rank test. **e,** Same as b, but with simultaneous optogenetic stimulation of pulvinar axons and optogenetic silencing of SOM cells. Left: the activity of V1 layer 2/3 cells was recorded while pulvinar axons and SOM cells were optogenetically co-manipulated for 3 s starting at grating stimulus onset (see methods). Second column: *n* = 256 cells, 5 sessions from 4 mice, *P* = 8.1 × 10^-4^, Wilcoxon signed-rank test. Third column: *n* = 256 cells, 5 sessions from 4 mice. Right: *n* = 40 and 41; *P* = 0.67 and *P* = 1.8 × 10^-4^; non-selective and highly selective cells; Wilcoxon signed-rank test. See also Extended Data Fig. 8 and 10.

Indeed, V1 neurons that responded to an unexpected stimulus (i.e. grating C) often also responded to the same stimulus when it was expected (Fig. 2a, b, c), but not to gratings A or B (Fig. 2a, b). Importantly, only neurons that responded highly selectively to a stimulus showed amplified responses when this stimulus was unexpected (Fig. 2d-f; *P* = 7.1 × 10^−4^ for highly selective cells). This selective amplification was also apparent in response to a second unexpected stimulus (grating D), or a familiar stimulus at an unexpected location (Extended Data Figs. 3 and 4). Moreover, the specific amplification of only selectively responding neurons could not be explained by differences in response strength (Extended Data Figs. 4g, h).

Together, these experiments indicate that the prediction error signal in layer 2/3 of V1 is an amplified response to the features of the unexpected visual input, rather than a non-specific surprise or a difference signal about how the visual input deviates from the animal’s predictions. Prediction errors specifically amplify the activity of neurons that respond highly selectively to the unexpected visual content, thereby selectively increasing the salience of unpredicted - and therefore potentially most relevant - sensory information.

### Prediction error signals require cortical disinhibition and pulvinar inputs

We next examined the circuit mechanisms by which sensory prediction errors are implemented in V1 networks. Vasoactive intestinal polypeptide (VIP)-expressing inhibitory interneurons in V1 receive cortical top-down and neuromodulatory inputs, and can disinhibit local principal cells through prominent inhibitory connections onto somatostatin (SOM)-expressing inhibitory interneurons^25–29^, providing a circuit for top-down gain modulation of sensory responses^30, 31^. VIP cells have also been shown to respond strongly to novel, but not familiar visual stimuli^21, 22^. To assess if VIP interneuron activity is important for prediction error signals in V1, we first examined how VIP interneurons respond to unexpected and expected visual information by using experimental paradigms described in Fig. 1k (Fig. 3a). VIP interneurons were suppressed by expected visual stimuli, but strongly responded to unexpected visual stimuli (Fig. 3b-d, Extended Data Fig. 5a-e), consistent with previous studies^15, 21, 22^. Responses of VIP interneurons decreased over time as mice encountered the same stimulus more often, in parallel with the gradual cessation of the prediction error signal in the layer 2/3 network (Fig. 3d, see also Fig. 1d), suggesting that the recruitment of VIP interneurons may be causally related to the generation of prediction error signals in V1.

**Fig. 5.**
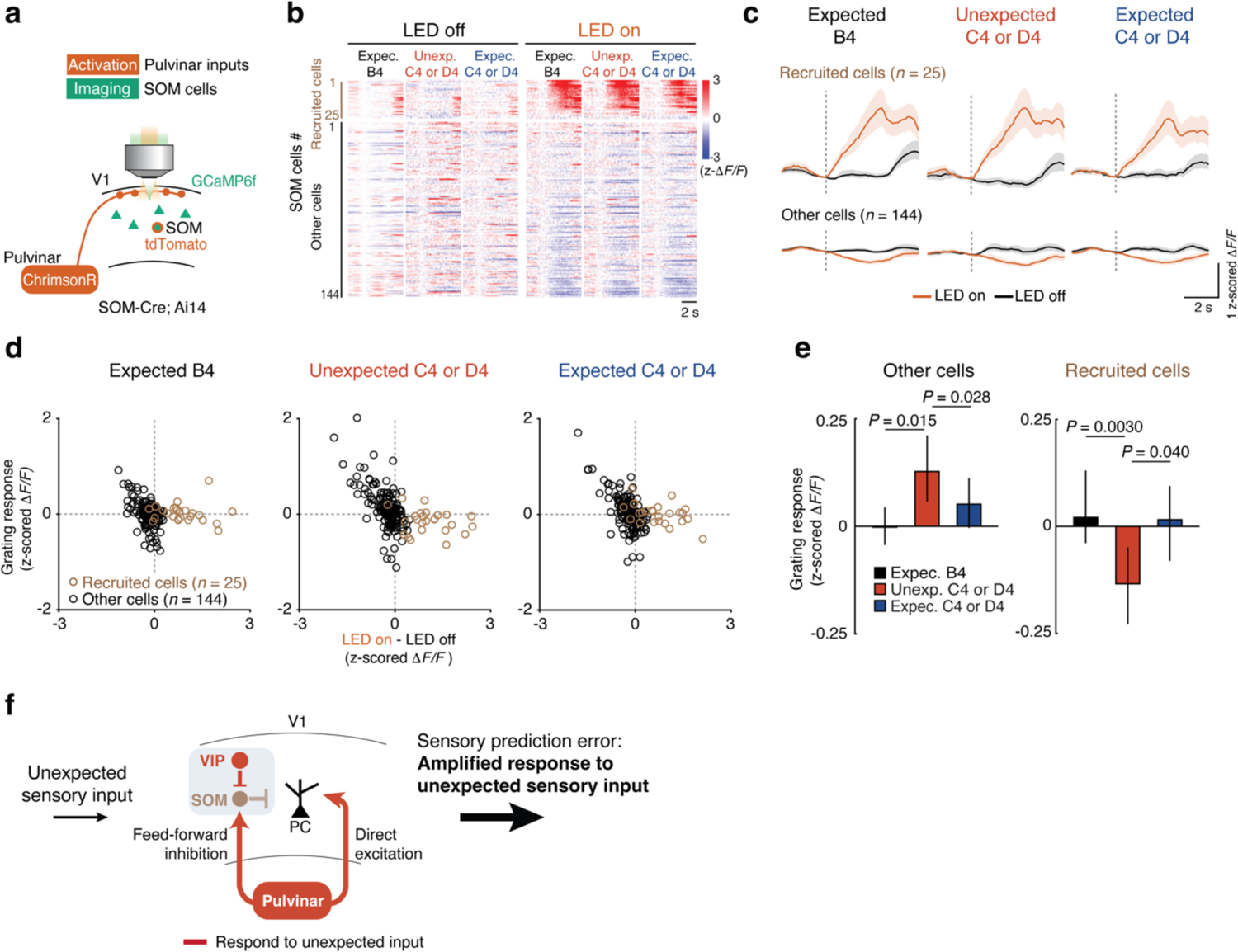
Pulvinar activates a specific subpopulation of SOM cells. **a**, Schematic of the experimental design. The activity of SOM cells was recorded while pulvinar axons were optogenetically stimulated for 3 s starting at grating stimulus onset. **b**, Single-cell responses for all SOM cells (individual rows, *n* = 169 cells, 5 sessions from 3 mice) to expected B4, unexpected C4 or D4 and expected C4 or D4 gratings with (right) or without (left) optogenetic stimulation. A subset of SOM cells was significantly recruited by optogenetic stimulation (recruited cells, *n* = 25; see Methods). **c,** Cell-averaged calcium responses with (amber) or without (black) optogenetic stimulation of recruited (*n* = 25) and other (*n* = 144) cells. Lines and shaded regions are mean and bootstrap 95% CI. **d**, Visual stimulus responses of individual SOM neurons to expected B4 stimulus (left), unexpected C4 or D4 stimulus (middle, in block 1) and expected C4 or D4 stimulus (right, in late block 2) plotted against the effect of pulvinar stimulation (difference in strength of visual stimulus responses with and without optogenetic pulvinar axon stimulation) for recruited (brown) and other (black) SOM cells. **e**, Strength of calcium response to expected B4 (black), unexpected C4 or D4 (red) and expected C4 or D4 (blue) stimuli of other SOM cells (left, B4 vs unexpected C4/D4: *P* = 0.015; unexpected vs expected C4/D4: P = 0.028, Wilcoxon signed-rank test with Bonferroni correction) and recruited SOM cells (right, B4 vs unexpected C4/D4: *P* = 0.0030; unexpected vs expected C4/D4: *P* = 0.040, Wilcoxon signed-rank test with Bonferroni correction). Bars and error bars indicate mean and 95% bootstrap CI. **f,** Proposed circuit mechanism for sensory prediction errors. VIP neurons inhibit a specific subpopulation of SOM cells that otherwise gate pulvinar input to V1, resulting in specific pulvinar-driven response amplification of the most stimulus-selective neurons in V1. See also Extended Data Fig. 9.

To test if the recruitment of VIP interneurons is required for the prediction error signal in the general V1 population, we optogenetically silenced VIP interneurons as mice encountered expected or unexpected visual stimuli while recording calcium responses of V1 layer 2/3 neurons (Fig. 3e-g, see Methods). Inactivating VIP neurons significantly reduced the responses of V1 layer 2/3 cells to unexpected visual stimuli (Fig. 3f middle; *P* < 1.0 × 10^−4^, Extended Data Fig. 5j, k), whereas it had no effect on responses to expected visual stimuli A and B (Fig. 3f left; *P* = 0.26), consistent with the specific recruitment of VIP interneurons by unexpected sensory stimuli (Fig. 3a-d). Furthermore, the effect of VIP inactivation on individual V1 layer 2/3 cells was not uniform, but highly correlated with how strongly they were facilitated by prediction errors: neurons with the strongest prediction error signal were the ones most suppressed by VIP interneuron inactivation (Fig. 3g). The effects of optogenetic inactivation on visually evoked responses could not be explained by light application artefacts, as light stimulation had no effect on responses to either unexpected and expected stimuli in control animals (Extended Data Fig. 6). Together, these experiments demonstrate that the recruitment of VIP interneurons is required for the generation of prediction error signals in layer 2/3 of V1.

Which long-range inputs to V1 could mediate the activation of VIP neurons by prediction errors? The pulvinar, a higher-order visual area in thalamus, integrates information from many cortical and subcortical areas and sends prominent feedback projections to V1^32–36^. Importantly, pulvinar projections to V1 carry information about visual input that is not predicted by the animal’s own actions, indicating that the pulvinar conveys sensory-motor prediction errors to V1^32^. To test if pulvinar projections to V1 also signal prediction errors arising from spatial predictions of visual input in our task, we used two-photon imaging to record calcium signals from pulvinar axons in V1^32^. Calcium activity of pulvinar axons was strongly and non-selectively boosted when a visual stimulus was unexpected (Fig. 3h-k, Extended Data Fig. 7), and this prediction error response decreased with repeated exposure to the same stimulus, with a time course similar to responses in V1 neurons (Fig. 3k).

To determine if pulvinar input to V1 is required for prediction error signals in V1 neurons, we optogenetically inactivated pulvinar axons in V1 while recording calcium responses of V1 layer 2/3 neurons (Fig. 3l-n). Suppressing pulvinar input to V1 specifically reduced the responses of V1 layer 2/3 neurons to unexpected visual stimuli (Fig. 3m middle, *P* < 1.0 × 10^−4^), but not to expected stimuli (Fig. 3m; *P* = 0.065 and *P* = 0.13 for visual stimuli A and B, and expected C and D, respectively). Similar to the effect of VIP neuron silencing, V1 neurons with strong prediction error responses were more likely to be strongly suppressed by pulvinar inactivation (Fig. 3n). Together, these cell type-specific inactivation experiments indicate that both intracortical VIP interneurons and pulvinar inputs contribute to prediction error signals in V1. Next, we investigated how these two circuit elements interact to generate the amplified responses to unexpected stimuli.

### Cooperative thalamocortical circuit mechanism for stimulus-selective response amplification

Pulvinar axons make synaptic connections onto VIP neurons in the neocortex^31^. A plausible scenario for how the pulvinar and neocortical VIP neurons interact to mediate prediction error signals may therefore involve pulvinar input activating VIP neurons in V1 which in turn boost pyramidal neuron responses to unexpected visual stimuli through the VIP-SOM disinhibitory circuit^25–27, 29^. To directly test this hypothesis, we optogenetically stimulated either pulvinar axons or VIP interneurons in V1 while monitoring neural responses of V1 layer 2/3 neurons to the expected grating stimuli in the virtual corridor (Fig. 4a-c, see also Extended Data Fig. 8). Consistent with a previous report^37^, stimulating pulvinar axons broadly suppressed responses to visual stimuli in V1 (Fig. 4b; *P* < 1.0 × 10^−4^). Moreover, stimulating pulvinar axons excited only a small subset of VIP neurons, and decreased VIP neuron prediction error responses (Extended Data Fig. 9). Optogenetically stimulating VIP interneurons had a minor effect on V1 activity, with a non-significant trend towards facilitating visual responses, unlike the strong amplification of stimulus-selective V1 neurons by prediction errors (Fig. 4c). Remarkably, simultaneous co-activation of both pulvinar axons and VIP neurons strongly facilitated visual responses of a subset of V1 neurons (Fig. 4d), indicating that pulvinar input and VIP neurons act synergistically, not additively. Moreover, response facilitation was specific to neurons that responded highly selectively to the visual stimulus that was paired with optogenetic stimulation, while non-selective neurons were suppressed, mimicking the prediction error signal in V1 (Fig. 4d, Extended Data Fig. 10a-d; compared to Fig. 2d, e). Our experimental evidence therefore does not support a direct pathway from pulvinar inputs onto VIP neurons to facilitate V1 responses, but pulvinar and VIP neurons are likely recruited independently, and act synergistically to provide stimulus-selective amplification of responses to unexpected stimuli in V1.

Our results indicate that when VIP neurons are activated, they can counteract the inhibitory influence pulvinar activation has on the V1 network. The main synaptic targets of VIP neurons are SOM interneurons that inhibit the apical dendrites of pyramidal cells^25–27, 29^. VIP neurons can therefore disinhibit pyramidal cells via the inhibition of SOM neurons. We hypothesized that pulvinar activation may recruit SOM neurons whose inhibitory influence on the V1 network may be alleviated when VIP neurons are simultaneously active. If this were the case, silencing SOM neurons while activating pulvinar should have effects similar to VIP neuron and pulvinar co-activation. Indeed, simultaneous optogenetic stimulation of pulvinar axons and inactivation of SOM neurons in V1 completely abolished the pulvinar-driven suppression of V1 activity (Fig. 4e; compared to Fig. 4b). Remarkably, this manipulation also strongly and specifically facilitated visual responses of V1 neurons responding highly selectively to the visual stimulus paired with the optogenetic manipulation, again mimicking the V1 prediction error signal (Fig. 4e), and suggesting that the pulvinar’s excitatory drive onto V1 pyramidal neurons is accompanied by a strong feed-forward inhibitory drive via SOM neurons.

While higher-order sensory thalamocortical pathways do not prominently target cortical SOM neurons^37–39^, at least a subset of SOM neurons in V1 has been shown to receive input from the pulvinar^31^. We imaged responses of V1 layer 2/3 SOM neurons while optogenetically stimulating pulvinar axons in V1, and found that while most SOM neurons were either not affected or even suppressed, a subset of SOM neurons (16 ± 11%; mean ± SD) was strongly activated by pulvinar stimulation (Fig 5a-c). Notably, SOM neurons that were recruited by pulvinar stimulation were suppressed by unexpected visual input, suggesting that this subset of SOM neurons is inhibited by VIP neurons (Fig. 5d, e). In contrast, layer 2/3 SOM neurons not recruited by pulvinar stimulation were activated by unexpected visual stimuli, similar to VIP neurons, suggesting that they do not receive strong inhibition from VIP neurons and/or are more strongly driven by the local excitatory layer 2/3 network (Fig. 5d, e), consistent with previous studies^29, 40, 41^. Together, these results show that excitatory drive from the pulvinar onto V1 pyramidal neurons is paralleled by a powerful inhibitory pathway via a specific subpopulation of SOM neurons. When VIP neurons are active simultaneously with pulvinar input they inhibit SOM neurons, thus reducing feed-forward inhibition from pulvinar to V1, and allowing pulvinar drive to strongly activate a subset of layer 2/3 pyramidal cells (Fig. 5f). These results therefore reveal a circuit mechanism for the generation of prediction error signals through synergistic interactions of pulvinar inputs and VIP neurons.

## Discussion

Here we describe a mechanism for boosting sensory reponses by prediction errors in V1 when animals’ expectations of visual stimuli at specific locations of a virtual environment are violated. Prediction errors selectively amplify the representation of unexpected visual input, via synergistic interactions of higher-order thalamic input and local VIP-SOM disinhibitory circuits in V1.

Prediction error responses are dependent on VIP neuron activity as well as input from the pulvinar, a higher-order visual nucleus in the thalamus that has previously been implicated in predictive processing, and conveys prediction error signals to V1^32, 33, 42^. Co-activation of pulvinar axons and VIP neurons in V1 can reproduce the selective amplification of V1 neurons even in the absence of prediction errors. Importantly, we found that pulvinar input to V1 is gated by VIP-SOM inhibitory interactions. The pulvinar suppresses the activity of V1 cells via a subpopulation of SOM neurons. To allow pulvinar input to amplify V1 responses, this inhibition has to be alleviated by activity in VIP neurons that inhibit SOM neuron responses (Fig. 5f). This mechanism may explain seemingly contradictory findings about how the pulvinar affects cortical activity^37, 43^ and establishes VIP neurons as a gate for higher-order thalamic input to V1. VIP neurons receive prominent neuromodulatory and top-down cortical input, and have been shown to be activated by salient events such as reward, punishment and novel stimuli^21, 22, 25, 27, 28, 30, 31, 44–46^. They can therefore regulate the pulvinar’s influence on visual processing in V1 depending on the relevance of visual stimuli or the animal’s behavioral state. Since VIP-SOM disinhibitory circuits and higher order thalamic feedback input are present throughout the cortical hierarchy^25–27, 29, 31, 35, 45^, this cooperative circuit mechanism may serve as a common computational motif in neocortical networks.

While VIP neurons and pulvinar inputs to V1 are broadly recruited by unexpected stimuli (Extended Data Figs 5a-i and 7), prediction error signals in V1 are observed only in subpopulations of neurons that are highly selective for the visual stimulus encountered. Our results point to a potential circuit mechanism for this selective response amplification in V1. We reproduced the selective amplification of only stimulus-selective V1 neurons by co-activating VIP neurons with pulvinar input to V1, but also when bypassing VIP activation by silencing SOM neurons while stimulating pulvinar input (Fig. 4d,e). Thus, selectivity of response amplification in V1 neurons does not depend on VIP neuron recruitment, but rather on pulvinar input more effectively driving V1 neurons with sharp tuning. This suggests a selective influence of pulvinar on subpopulations of stimulus-selective V1 neurons, balanced by inhibition from pulvinar-driven SOM neurons (Extended Data Fig. 10e-h). This pulvinar-dependent response enhancement may be further amplified via recurrent excitation within subnetworks of selective V1 neurons tuned to the same stimulus^47^ and lateral suppression of the rest of the network via PV neurons^48–50^, collectively leading to selective amplification of unexpected input.

We observed sensory prediction error signals not only in V1, but also in the pulvinar. So where do prediction error signals arise? They may be computed downstream of these visual areas and conveyed to V1 by top-down projections via pulvinar and local VIP interneurons. Alternatively, errors could be computed in the pulvinar or in V1 from the comparison of visual input with predictions conveyed by top-down input from higher-order cortical areas^5–10, 14^, and amplified through pulvinar-V1 recurrent connections. Either way, signals in V1 may be further enhanced by neuromodulators such as acetylcholine or noradrenaline^28, 46^ that may signal stimulus saliency and novelty, or surprise more generally^51, 52^.

Our results indicate that individual V1 neurons do not signal how the actual visual input deviates from the animal’s predictions, as postulated within the predictive coding framework^5–8^. Instead, we propose an alternative view of predictive processing in sensory circuits: prediction errors amplify the representation of feedforward sensory input in neocortex, while the extent of amplification may depend on how much the visual stimulus deviates from expectations and therefore the magnitude of animals’ surprise. This would explain the particularly strong responses to novel stimuli that were not encountered before, as these are least expected^21, 22^. The amplified responses to unexpected stimuli may serve as a neural substrate for attentional shifts towards surprising events in the environment. However, the content of how actual input deviates from predictions may still be encoded in other brain areas or higher-dimensional population activity in V1.

In summary, sensory prediction errors in V1 increase the saliency of unexpected, and thus likely relevant visual information. This enables downstream brain areas to prioritize these signals and potentially utilize them for updating internal predictions.

## Methods

### Mice

All experiments were performed under the UK Animals (Scientific Procedures) Act of 1986 (PPL PD867676F) following local ethical approval by the Sainsbury Wellcome Centre Animal Welfare Ethical Review Body. A total of 76 mice, including 19 C57BL/6J mice, 24 VIP-Cre mice (JAX 010908, Jackson Laboratory; Cre expressed in VIP interneurons), 23 VIP-Cre × Ai14 mice (JAX 010908 and JAX 007914, Jackson Laboratory; tdTomato expressed in VIP interneurons), 7 SOM-Cre mice (JAX 013044, Jackson Laboratory; Cre expressed in SOM interneurons) and 3 SOM-Cre × Ai14 mice (JAX 013044 and JAX 007914, Jackson Laboratory; tdTomato expressed in SOM interneurons) were used in this study. Both female and male mice, at least 7 weeks old at the start of the experiments, were used. Mice were co-housed with littermates in IVC cages, in reversed day–night cycle lighting conditions, with the ambient temperature and humidity set to 23 °C and 56% relative humidity, respectively. Standard environment enrichment was provided in the form of a running wheel, a clear tube and wooden toys.

### Surgical procedures

Prior to surgery, Dexadreson (2-3 mg per kg) and Carprofen (5 mg per kg) were administered. General anesthesia was induced with 2.5-3% isoflurane, which was then reduced to maintain a breathing rate of around 1 Hz. A 3 or 4 mm craniotomy was made over the right V1, centered on 2.45 mm lateral and 3.6 mm posterior of bregma. For two-photon calcium imaging and optogenetic manipulations of V1 cells, we injected AAV vectors into right monocular V1 (centered on 2.45 mm lateral and 3.7 mm posterior of bregma, 1-3 injections per mouse, 100-150 nL per injection). For two-photon calcium imaging and optogenetic manipulations of pulvinar axons, we injected AAV vector into right pulvinar (calcium imaging and optogenetic activation: 1.6 mm lateral and 2.1 mm posterior of bregma, 2.35 below the cortical surface, 1 injection per mouse, 60 nL per injection; optogenetic inactivation: 1.55 mm lateral and 2.0 mm posterior of bregma, 2.3 mm below the cortical surface, 1.60 mm lateral and 2.2 mm posterior of bregma, 2.4 mm below the cortical surface, 2 injections per mouse, 60 nL per injection). All injections were performed using glass pipettes and Nanoject III microinjector (Drummond Scientific) or a pressure injection system (Picospritzer III, Parker). A 3 or 4 mm circular cover glass was glued in place using cyanoacrylate glue (Pattex). A custom-designed stainless steel headplate was attached to the skull using dental cement (Super-Bond C&B, Sun Medical). Animals were given analgesics (Carprofen; 5 mg/kg) at 24 and 48 hours after surgery. Imaging started approximately 3 weeks after the virus injection.

### Viral constructs

We used AAV1-hSyn-GCaMP6f (2 × 10^13^ vg/mL Penn Vector Core/Addgene; diluted 1:8 to 1:15 in saline) for experiments involving two-photon calcium imaging of V1 layer 2/3 cells; AAV1-hSyn-GCaMP7b (2 × 10^13^ vg/mL Penn Vector Core/Addgene; diluted 1:2 in saline) for imaging of pulvinar axons; AAV2-EF1a-DIO-eNpHR3.0-mCherry (4.0 × 10^12^ vg/mL, 1:2 to 1:10 dilution, UNC vector core) for optogenetic silencing of VIP cells or SOM cells; AAV2-hSyn-eNpHR3.0-mCherry (3.3 × 10^12^ vg/mL, 1:2 to 1:4 dilution, UNC vector core) for optogenetic silencing of pulvinar axons; AAV1-hSyn-Flex-ChrimsonR-tdTomato (3.9 × 10^12^ vg/mL, 1:2 to 1:5 dilution, UNC vector core) for optogenetic activation of VIP cells; AAV1-Syn-ChrimsonR-tdTomato (4.1 × 10^12^ vg/mL, 1:2 to 1:5 dilution, UNC vector core) for optogenetic activation of pulvinar inputs; AAV1-hEF1a-mCherry (5.7 × 10^12^ vg/mL, 1:2 to 1:5 dilution, Zurich vector core) for control experiment for LED light stimulation.

### Behavioral setup

Behavioral setups consisted of a styrofoam running wheel, two visual stimulation display monitors (see below), a reward delivery spout, and a camera for recording the pupil. Mice were head-fixed and placed on a styrofoam wheel (20 cm diameter, 12 cm width). Their running speed was monitored using a rotary encoder (Kubler Encoder 1000 ppr) coupled to the wheel axle. Reward (a drop of strawberry milk, 50% Ensure nutrition shake, Abbott Laboratories) was delivered by a lick spout in front of the mouse and was regulated via a solenoid pinch valve (161P011, NResearch). Licks were detected with a piezoelectric diaphragm sensor (7BB-12-9, Murata) placed under the spout. Images of the left eye were recorded with a CMOS camera (22BUC03, Imaging Source) at 30 Hz in order to track eye movements and pupil size. The recording of the encoder, presentation of visual stimuli, opening of the reward valves, and camera recordings were controlled by custom-written software in LabView. Behavioral data were acquired using a PCIe 6321 acquisition card (National Instruments).

### Food restriction and pre-training

Before animals underwent training in the virtual environment, they were food restricted and pre-trained to encourage continuous running on the styrofoam wheel. Four to seven days after surgery, food-restriction and pre-training started. Mice were weighed daily and given typically 2-3 g of food pellet in addition to strawberry milk given in training sessions to ensure they maintain around 90%, but at least 85% of their starting body weight. For the first few days, animals were handled in a soft cloth and iteratively fed strawberry milk (Abbott Laboratories) through a syringe until they got used to short manual restraint of the headplate. Mice were then head-fixed and put on the freely rotating styrofoam wheel for 15 to 60 min. Mice were encouraged to run on the wheel by delivering strawberry milk rewards after they moved a short distance (initially set to ∼10 cm). This distance was adjusted (up to 500 cm) depending on the running speed of the mouse, such that mice received roughly one reward every 30 s. Additional rewards were occasionally delivered by the experimenter. This pre-training took 4-10 days.

### Virtual corridor

Once mice were running continuously, they were moved to a virtual environment consisting of a linear corridor with varying wall patterns as described previously^14^. The cylinder’s rotation (the instantaneous running speed of the animal) was used to control the speed at which the animal moved through the virtual environment. The virtual environment was displayed on two monitors (U2715H, Dell; 60 Hz refresh rate), placed 21 cm away from both eyes of mice and oriented at 35° relative to the midline. Each monitor covered a visual field of approximately 110 degrees horizontally and 60 degrees vertically. All elements of the corridor including the gratings were calibrated to be isoluminant (10.1 cd/m^2^). The luminance of the monitor was set at 0.1 cd/m^2^, 10.1 cd/m^2^ and 20.1 cd/m^2^, at black, gray, and white values, respectively. The luminance of visual stimuli was measured using a luminance meter (Konica Minolta, LS-100). The gray walls of the virtual corridor were lined with four different landmarks. The last landmark represented the reward zone located at the end of the corridor. Reaching the reward zone triggered an automatic reward delivered by a spout located in front of the mouse. After the reward delivery, the virtual environment was reset to the beginning of the corridor to start the next trial.

Grating stimuli were suddenly presented on full screen once the mouse entered a certain position in the corridor. This was done to ensure precise control of when the mouse would first see the grating. Grating stimuli were presented at four different positions between landmarks. The optic flow of the gratings was ‘uncoupled’ from the running speed for 2.4 s, such that the animal’s locomotion did not affect its temporal frequency. Gratings were square-wave gratings, with the spatial frequency of approximately 0.04 cycles per degree (cpd) at the center of the monitor and the temporal frequency approximately 2 cycles per second (Hz). Duration of the grating presentation was approximately 2 s at the center of the monitors. The precise timing of visual stimulus onsets was recorded with a photodiode (Thorlabs) attached to the monitor.

During 5 training sessions, the virtual corridor and the sequence of the four grating stimuli was identical (A-B-A-B) on every trial. In the subsequent imaging session, the identity of one of the four gratings was changed. In block 1 of this session (160 trials), the identity of the 4^th^ grating stimulus B changed either to a novel stimulus grating C (C session), a novel stimulus D (D session), familiar stimulus A (A session) or no grating was shown (omission session) on randomly chosen 10 % of trials. In block 2, the novel stimulus or no stimulus was shown at the fourth position in 100 % of trials. Occasionally one mouse underwent several sessions with unexpected stimuli. In that case, mice went through another training session (with gratings A-B-A-B) in between. For imaging of pulvinar axons, block 1 was shortened to 60 trials, and either a novel stimulus grating C (C session) or a novel stimulus D (D session) was shown at the fourth position on randomly chosen 15 % of trials. In block 2, the novel stimulus was shown in 100 % of trials, as for imaging of V1 layer 2/3 cells.

### Two-photon calcium imaging

Two-photon calcium imaging was performed using a commercial resonance scanning two-photon microscope (B-Scope; Thorlabs) with a 16× water-immersion objective (NA 0.8, Nikon), with a Ti::Sapphire laser at 930 nm excitation wavelength (Mai Tai, SpectraPhysics). Emission light was band-pass filtered using a 525/50 filter (Semrock) or a 520/40 filter (Chroma) for GCaMP, and a 607/70 filter for tdTomato/mCherry (Semrock). Images of 512 × 512 pixels from four imaging planes with fields of view ranging from 380 ×380 μm to 440 × 440 μm were acquired at 7.5 Hz frame rate for imaging of V1 neurons and of a single plane of 160 × 160 μm at 15 Hz frame rate for imaging of pulvinar axonal boutons using ScanImage^53^. For imaging of V1 neurons, we used a piezo-actuator (Physik Instrumente) to move the objective in steps of 15 μm between frames to acquire images at four different depths, thus reducing the effective frame rate to 7.5 Hz. Imaging of V1 neurons was performed in layer 2/3 (typically 150-200 μm below the cortical surface). The laser power under the objective never exceeded 35 mW. Axonal bouton calcium measurements were performed in cortical layer 1 (35-55 μm below the cortical surface), with laser powers below 20 mW.

To avoid cross-talk between imaging and visual stimulation, the monitor backlight was controlled using a custom-built circuit to present visual stimuli only at the resonant scanner turnaround points in between two subsequent imaging lines (when data were not acquired)^54^. The frame trigger signal during two-photon calcium imaging was recorded by Labview and used for synchronization between the calcium imaging frames and task related data (e.g., behavior data and visual stimuli onsets).

For imaging of pulvinar axons, we used VIP-Cre × Ai14 mice. We simultaneously imaged pulvinar axons expressing GCaMP7b and neurites of VIP neurons expressing tdTomato in layer 1. We then used the red signal (tdTomato) as a structure marker to perform Z-drift correction during imaging and frame registration in data preprocessing.

### Optogenetic manipulation

Simultaneous two-photon imaging and optogenetic stimulations were performed as previously described^15^. Briefly, 595 nm light was delivered through the objective lens using a fast LED (UHP-T-595, Prizmatix). The LED light power was set to 8 mW in front of the objective. To combine two-photon imaging and optogenetic manipulation, the LED for optogenetic manipulation was synchronized to the resonant scanner turnaround points (when data were not acquired). The propagation of reflected light to the eyes of the mouse was blocked by a metal light shield cone placed on the headplate and a black cement wall around the implant. Optogenetic manipulation occurred in randomly chosen 10 – 50 % of each trial type. Two protocols for LED application were employed in this study: continuous stimulation for 3 seconds starting at grating stimulus onset (Figs 3, 4e, 5, Extended Data Figs. 5k, 6, 8d-e, 9, and 10e-h), or stimulation for 1 second at a frequency of 20 Hz, with 40% duty cycle (20 ms pulses), starting 0.1 second after grating stimulus onset (Fig 4 b-d and Extended Data Figs. 8a-c, 10a-d).

### Histology

At the end of each experiment, targeting of virus injections was confirmed by histology. Mice were euthanized with a dose of pentobarbital (80 mg/kg) and transcardially perfused with 4% paraformaldehyde. Brains were extracted and post-fixed overnight in 4% paraformaldehyde, stored in a 50 mM phosphate buffer. Brains were embedded in 5% agarose and imaged using a custom-built serial-section two-photon microscope. Coronal slices were cut at a thickness of 50 μm using a vibratome (Leica VT1000), and imaged at two optical planes per physical section. Scanning and image acquisition were controlled by ScanImage v5.6 (Vidrio Technologies, USA) using a custom software wrapper for defining the imaging parameter. A subset of brains was embedded in 4% agarose (A9539, Sigma), cut in 200 μm coronal slices on a vibratome (HM650V; Microm), mounted in a mounting medium containing DAPI (Vectashield; Vector Laboratories) and imaged on a slide scanner (Zeiss AxioScan) or on a confocal microscope (Leica SP8).

### Quantification and statistical analysis

#### Two-photon imaging

Two-photon imaging frames were motion corrected and segmented using custom-written scripts in MATLAB as previously described^32^. Briefly, to correct for x-y motion, two-photon imaging frames were registered to a 1200-frame average (40-frame × 30 batches) using a phase-correlation algorithm. In Fig. 1l-n and Extended Data Fig. 5b-i, images from C session and D session were registered together, and identical cells were matched across sessions by using custom-written software. Frames with large motion were detected by inspecting the registration displacement results and were discarded from further analysis. Regions of interest (ROIs) were detected semi-automatically using intensity thresholding combined with PCA-ICA refinement and validated and refined manually. All time-series were extracted and analyzed with custom written functions using the TimeSeriesAnalysis package^55^. All pixels within each ROI were averaged to give a single time course. Contaminating signals from neuropil were subtracted using an Asymmetric Student-t model (ast_model)^56^. Calcium *ΔF/F_0_* signals were obtained by using the baseline fluorescence *F*_0_, which is estimated by a gaussian mixture model with two components fitted on the raw fluorescence data. The mean parameter of the lowest Gaussian component is used as *F*_0_. To be able to compare calcium activity across sessions and mice, z-scored *ΔF/F* was computed by subtracting the mean value of *ΔF/F* of a session and dividing the resulting trace by the standard deviation.

#### Analysis of visual responses

The response to each grating was calculated using the mean z-scored *ΔF/F* calcium signal averaged over a window from 0.4 s to 2 s after grating onset, baseline-subtracted using the mean z-scored *ΔF/F* signal during 0.5 s before stimulus onset for each grating presentation. Neurons were classified as stimulus-responsive if their mean response was bigger than 0.5 z-scored *ΔF/F*. In Fig. 5 and Extended Data Fig. 9, a longer time window from 0.4 s to 3 s after grating onset was used as response window.

In Fig. 1, cell-averaged calcium traces are from neurons significantly responsive to the presented grating in unexpected C or D trials (block 1), expected C or D trials (late block 2) or both. Similar average response traces, but including all neurons responsive to any grating are shown in Extended Data Fig. 2. In Fig. 2 a-b and Extended Data Fig. 4 a-b, cells were defined as prediction-error responsive if the responses were significantly different between unexpected C4 or D4 (block 1) and expected C4 or D4 (second half of block 2, two-sided t-test; α = 0.05; unexpected C4 or D4 versus expected C4 or D4) and the difference in response was larger than 0.5 z-scored *ΔF/F*. Similarly, in Extended Data Fig. 5 e and i, cells were defined as prediction-error responsive if the responses were significantly different between unexpected C4 or D4 (block 1) and expected C4 or D4 (second half of block 2, two-sided t-test; α = 0.05; unexpected C4 or D4 versus expected C4 or D4) and the difference in response was larger than 0.3 z-scored *ΔF/F*. In Fig. 5, SOM cells were defined as ‘recruited’ if their responses were significantly different with and without optogenetic pulvinar axon stimulation (two-sided t-test; α = 0.016; with versus without LED light stimulation) and the difference in response was larger than 0.3 z-scored *ΔF/F*, during at least one of the visual stimuli presentations (expected B4, unexpected C4 or D4, expected C4 or D4). In Extended Data Fig. 9, VIP cells were defined as ‘recruited’ if the responses to expected grating B4 were significantly different with and without optogenetic stimulation (two-sided t-test; α = 0.05) and the difference in response was larger than 0.3 z-scored *ΔF/F*. In Fig. 3 f, g, m and n, neurons were included if their average response to the visual stimulus presented with and without LED stimulation was larger than 0.5 z-scored *ΔF/F,* in order to avoid selection bias in the response strength towards LED off trials. In Fig. 4, neurons were defined as stimulus responsive if their stimulus response strength was larger than 0.5 z-scored *ΔF/F* in trials without LED stimulation in order to avoid signal contribution of opsin-expressing and therefore directly activated VIP cells.

#### Selectivity and selectivity index

To quantify the selectivity of neural responses we computed a response selectivity measure for individual V1 layer 2/3 cells:

Selectivity = (*R*_C4 or D4_ - *R*_A3 or B2_) / (*R*_C4 or D4_ + *R*_A3 or B2_)

Where *R*_C4 or D4_ is the mean response to the gratings C4 or D4 in late block 2, and *R*_A3 or B2_ is the mean response to the gratings A3 or B2 in late block 2. Selectivity values of > 1 or < −1 were shown as 1 or −1, respectively. If selectivity was less than 0.6 or more than 0.8, they were classified as either non-selective or highly selective, respectively.

We used a different selectivity index (SI) to quantify response selectivity of individual pulvinar boutons (Extended Data Fig. 7), since this index provided a more reliable measure for the noisy bouton calcium traces. SI was calculated as previously described^57^. Briefly, it was computed from the difference between the mean response to the expected stimulus C4 or D4 and expected stimulus A3 in late block 2, divided by the pooled standard deviation of the responses.

#### Fast and slow running trials

For the analysis in Extended Data Fig. 1, trials in block 2 were divided into fast and slow running trials based on mean running speed during presentation of grating C4. A time window starting 0.4 s and ending 2 s after the grating onset, similar to the response window for *ΔF/F*, was used to calculate the mean running speed. A trial was defined as ‘fast’ or ‘slow’ if the mean running speed during the time window was in the top 40th or bottom 40th percentile of all visual stimuli C4 presentations in block 2.

#### Correlation of running speed and neuronal activity

To determine the effect of running speed on neuronal activity, we computed for each cell the correlation between mean *ΔF/F* and mean running speed in a time window (starting 0.4 s and ending 2 s after grating stimulus onset) of each trial in block 2. We used the square of the correlation coefficient (R^2^, coefficient of determination) to quantify the strength of the modulation of neural responses by running speed.

#### Pupil size

Pupil size was computed offline. Pupil was detected and determined using a binary threshold and center of mass of the detected regions. We then applied a one-dimensional filter to the traces using *filloutlier* function in MATLAB.

#### Statistics

We used two-sided Wilcoxon rank-sum tests for independent group comparisons, and two-sided Wilcoxon signed-rank tests for paired tests, unless otherwise stated. Raw p-values, except for those that are below 1.0 × 10^−4^, are reported throughout the manuscript. Significance thresholds were adjusted for multiple comparisons using Bonferroni correction, as indicated in the figure legends. Tests were performed using MATLAB. The mean and the bootstrap 95% confidence intervals were used for display purposes, as stated in the figure legends. Confidence intervals were estimated using *bootci* function in MATLAB, with 10,000 bootstrap samples with replacement. Linear regression models were fitted using the *fitlm* function in MATLAB. No statistical methods were used to predetermine sample sizes, but our sample sizes are similar to those generally employed in the field.

## Acknowledgements

We thank M. Li for help with animal husbandry and pre-training; M. Rio for help with calcium data preprocessing pipeline and virtual reality; R.A.A. Campbell for help with microscopy; T. Kanamori, I. Voitov and M. Javadzadeh for feedback on the manuscript; the T.D.M.F. laboratory and the S.B.H. laboratory for discussions. This work was supported by a Sainsbury Wellcome Centre Core Grant from the Gatsby Charitable Foundation and Wellcome (219627/Z/19/Z and 090843/F/09/Z); a Wellcome Investigator Award (219561/Z/19/Z to S.B.H.); the Gatsby Charitable Foundation (GAT3212 and GAT3361 to T.D.M.-F.); the Wellcome Trust (090843/E/09/Z and 217211/Z/19/Z to T.D.M.-F.); European Research Council (HigherVision 337797 to S.B.H; NeuroV1sion 616509 to T.D.M.-F); the SNSF (31003A 169525 to S.B.H.); Biozentrum core funds (University of Basel).

## Author contributions

S.F., S.B.H. and T.D.M.-F. conceived the study. S.F. performed the experiments and analyzed the data. A.D.F. assisted with surgical procedures, animal pre-training and preliminary optogenetic experiments. S.F., S.B.H., and T.D.M.-F. wrote the manuscript.

## Data and code availability

The data reported in this study and analysis code are available from the corresponding authors upon request.

---

**Extended Data Fig. 1.**
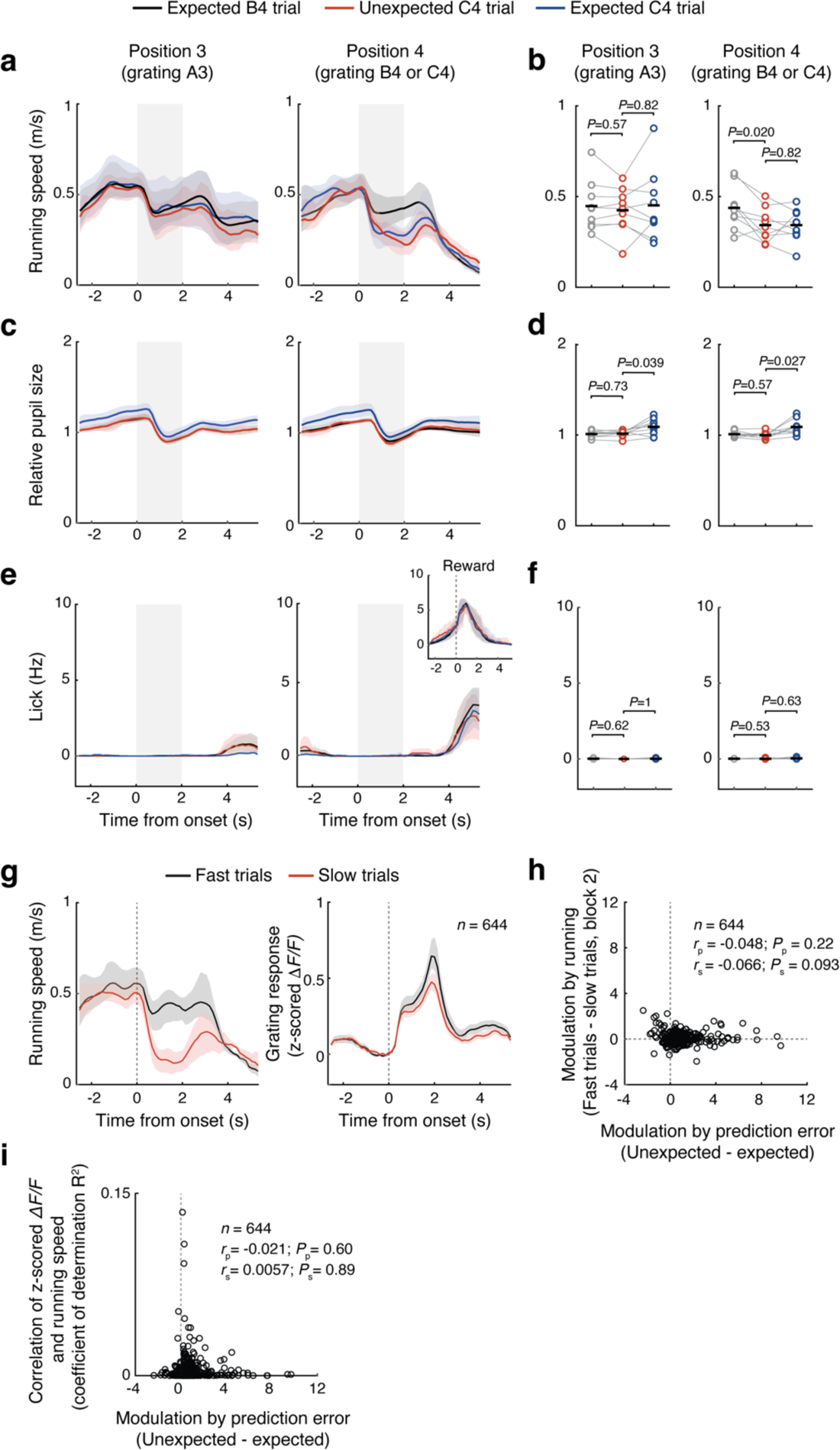
Running and licking behavior and pupil size during presentation of expected and unexpected gratings. Related to Figs. 1 and 2. **a**, Running speed at virtual corridor position 3 (left, grating A3 shown) or at position 4 (right, grating B4 or C4 shown) in trials in which grating B was presented at position 4 (black, expected B4 trials, 90% of trials in block 1), trials in which grating C was presented at position 4 (red, 10% of trials in block 1, unexpected C4 trials) and trials in the second half of block 2 (blue, expected C4 trials, late block 2). Light gray shading indicates length of visual stimulus at the center of monitors. Lines and shading are mean and bootstrap 95% CI (*n* = 9 mice). **b**, Same as a, but data from individual animals are shown separately. Black bars represent mean across animals. Position 3, running speed during grating A3 presentation in B4 vs unexpected C4 trials: *P* = 0.57; running speed during grating A3 presentation in unexpected vs expected C4 trials: *P* = 0.82. Position 4, running speed during B4 vs unexpected C4 presentation: *P* = 0.020; running speed during unexpected vs expected C4 presentation: *P* = 0.82. *n* = 9 mice, Wilcoxon signed-rank test with Bonferroni correction. **c** and **d**, Same as a and b, but for relative pupil size (normalized by each session’s median value). **d**, Position 3, pupil size during grating A3 presentation in B4 vs unexpected C4 trails: *P* = 0.73; pupil size during grating A3 presentation in unexpected vs expected C4 trials: *P* = 0.039. Position 4, pupil size during B4 vs unexpected C4 presentation: *P* = 0.57; pupil size during unexpected vs expected C4 presentation: *P* = 0.027; *n* = 9 mice, Wilcoxon signed-rank test with Bonferroni correction. **e** and **f**, Same as a and b, but for lick rate. **e**, Inset shows lick rate around the reward delivery. **d**, position 3, lick rate during A3 presentation in B4 vs unexpected C4 trials: *P* = 0.62; lick rate during A3 presentation in unexpected vs expected C4 trials: *P* = 1. Position 4, lick rate during B4 vs unexpected C4 presentation: *P* = 0.53; lick rate during unexpected vs expected C4 presentation: *P* = 0.63. *n* = 9 mice, Wilcoxon signed-rank test with Bonferroni correction. **g**, Running speed (left) and responses to grating C4 (right) on trials of fast (black, in the top 40%) and slow (red, in the bottom 40%) running speed during grating C4 presentation at position 4 in block 2 (see Methods). **h**, Scatterplot showing no correlation between response modulations by running speed and strength of prediction error responses (Pearson correlation: *r*_p_ = −0.048, *P*_p_ = 0.22; Spearman correlation: *r*_s_ = −0.066, *P*_s_ = 0.093; *n* = 644 cells from 9 mice). **i**, Scatterplot showing no correlation between correlation of z-scored *ΔF/F* and running speed (coefficient of determination R^2^) and strength of prediction error responses (Pearson correlation: *r*_p_ = −0.021, *P*_p_ = 0.60; Spearman correlation: *r*_s_ = 0.0057, *P*_s_ = 0.89; *n* = 644 cells from 9 mice).

**Extended Data Fig. 2.**
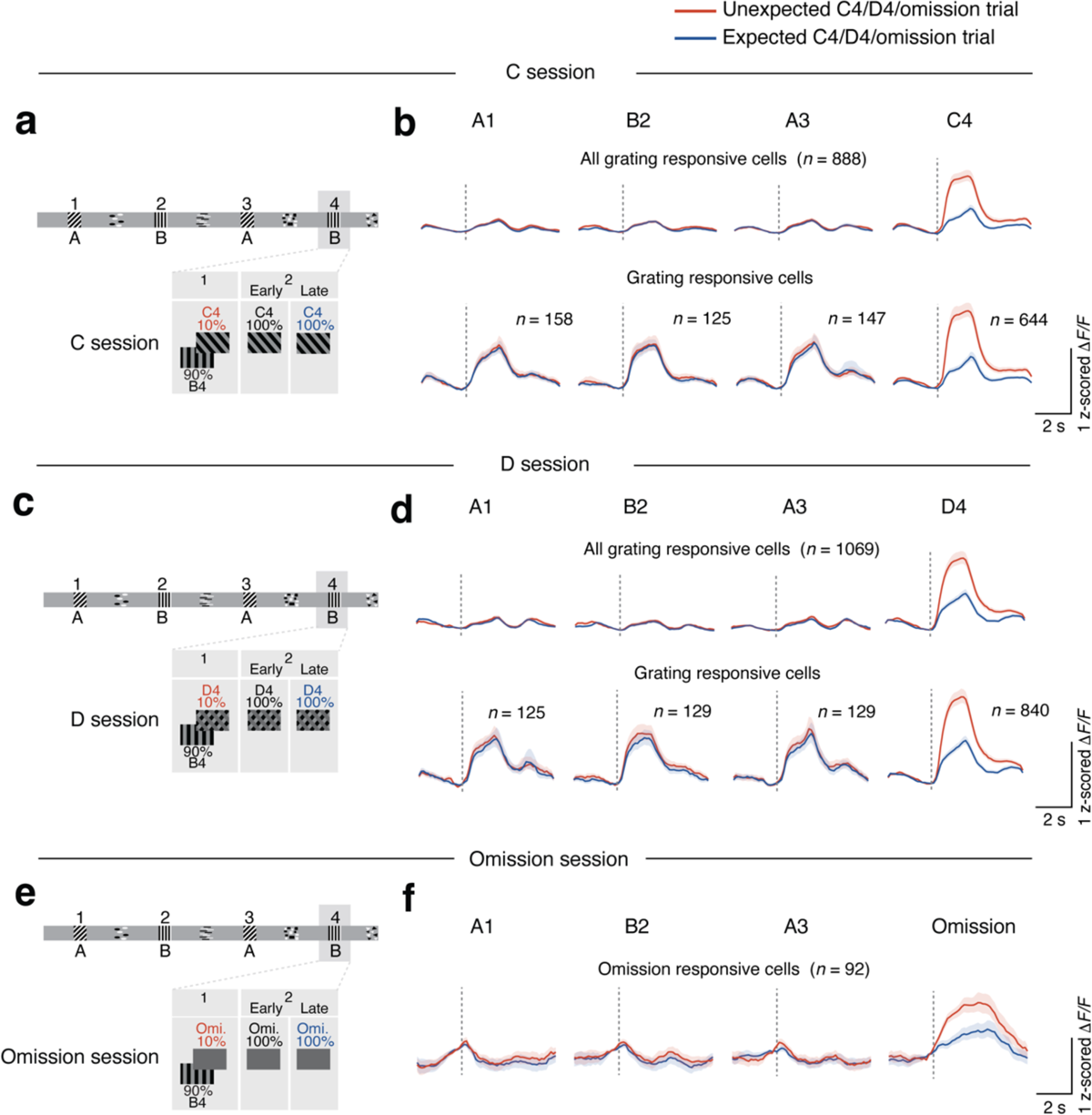
Average calcium responses in V1 to expected and unexpected stimuli and unexpected stimulus omissions. **a**, Schematic of visual stimuli shown in a C session (unexpected C stimulus presented in 10% of trials instead of B at position 4 in block 1 and in 100% of trials in block 2). **b**, Top: Average calcium responses of all cells significantly responsive to any of presented gratings in unexpected C4 (block 1) or expected C4 (late block 2) trials (*n* = 888). Dotted line indicates grating stimulus onset. Bottom: Average calcium responses as on top, but only of neurons significantly responsive to each presented grating (*n* = 158, 125, 147, 644; Gratings A1, B2, A3, C4 responsive cells). Same as Fig. 1c. Data from 9 mice. **c**, Schematic of visual stimuli shown in a D session (unexpected D stimulus presented in 10% of trials instead of B at position 4 in block 1 and in 100% of trials in block 2). **d**, Same as **b** but responses to D4 on the right. Top: *n* = 1069. Bottom: *n* = 125, 129, 129, 840; A1, B2, A3, D4 responsive cells. Data from 7 mice. **e**, Schematic of visual stimuli shown in an omission session (stimulus B4 omitted in 10% of trials in block 1 and in 100% of trials in block 2). **f**, Average calcium responses to gratings A1, B2, A3 and omission of B4 of all omission responsive cells (*n* = 92 from 5 mice). Lines and shading are mean and bootstrap 95% CI.

**Extended Data Fig. 3.**
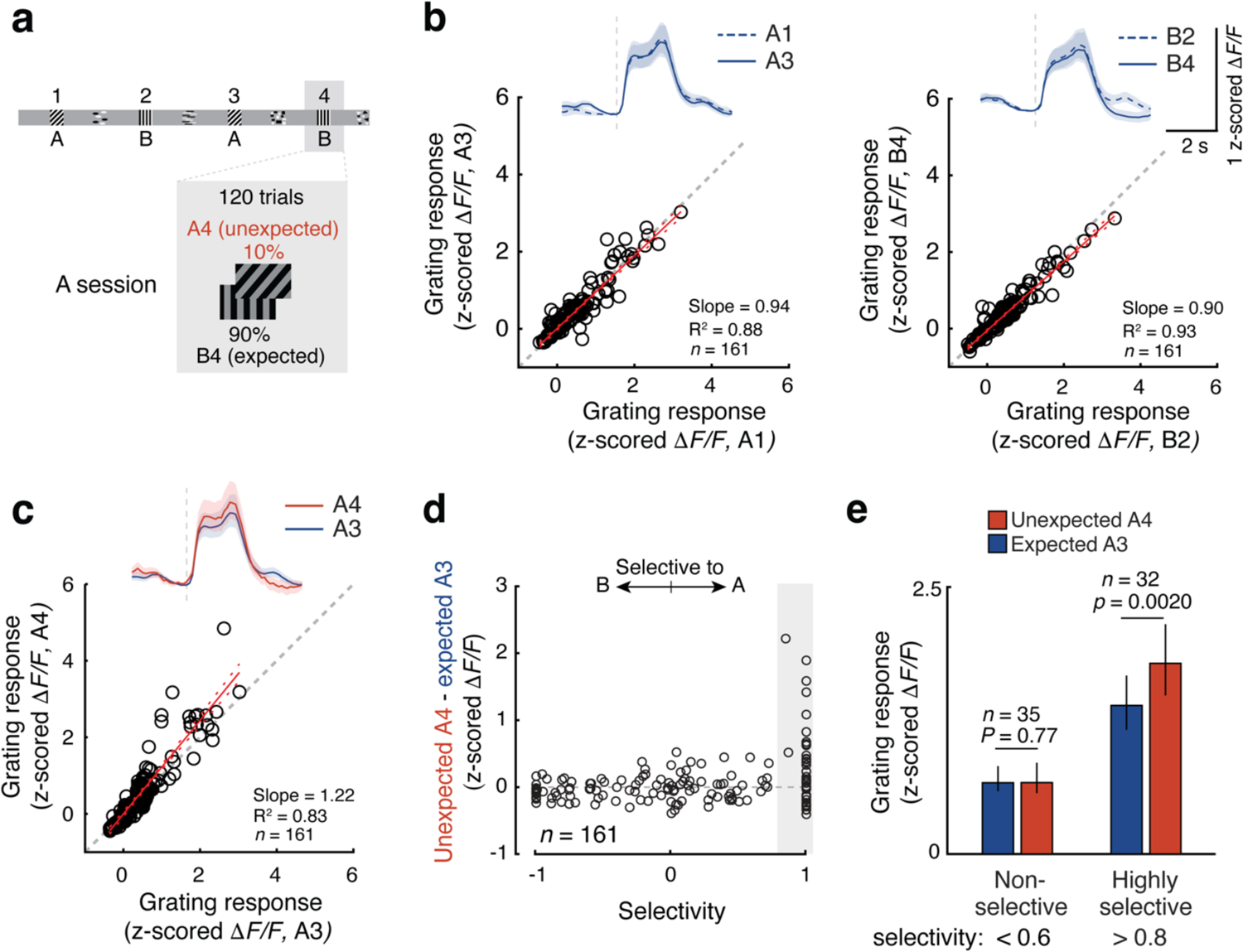
Prediction error in response to familiar visual stimulus encountered at an unexpected location (grating A presented at location 4 instead of grating B). **a**, Schematic of experimental design. **b**, Left: calcium responses of individual V1 neurons to expected grating A1 plotted against responses to expected grating A3. Right: calcium responses of individual V1 neurons to expected grating B2 plotted against responses to expected grating B4. Neurons responsive to at least one of the expected grating stimuli (A1, B2, A3 or B4) were included in the analysis. *n* = 161 cells from 8 mice. R^2^: coefficient of determination. Red dotted lines: 95% confidence interval of the linear fit. **c**, Calcium responses of individual V1 neurons to expected grating A3 plotted against responses to unexpected grating A4. R^2^: coefficient of determination. Red dotted lines: 95% confidence interval of the linear fit. **d**, Strength of prediction error signal (difference in response to unexpected and expected grating A) plotted against grating response selectivity (difference in response to grating A and grating B divided by the sum of responses) for all grating-responsive cells. **e**, Cell-averaged response strength to expected grating A3 (blue) and unexpected grating A4 (red) of non-selective (selectivity A3 vs B2 < 0.6, left, *n* = 35, *P* = 0.77, Wilcoxon signed-rank test) and highly selective (selectivity A3 vs B2 > 0.8, right, *n* = 32, *P* = 0.0020) grating A3 responsive cells. Bars and error bars are mean and 95% bootstrap CI.

**Extended Data Fig. 4.**
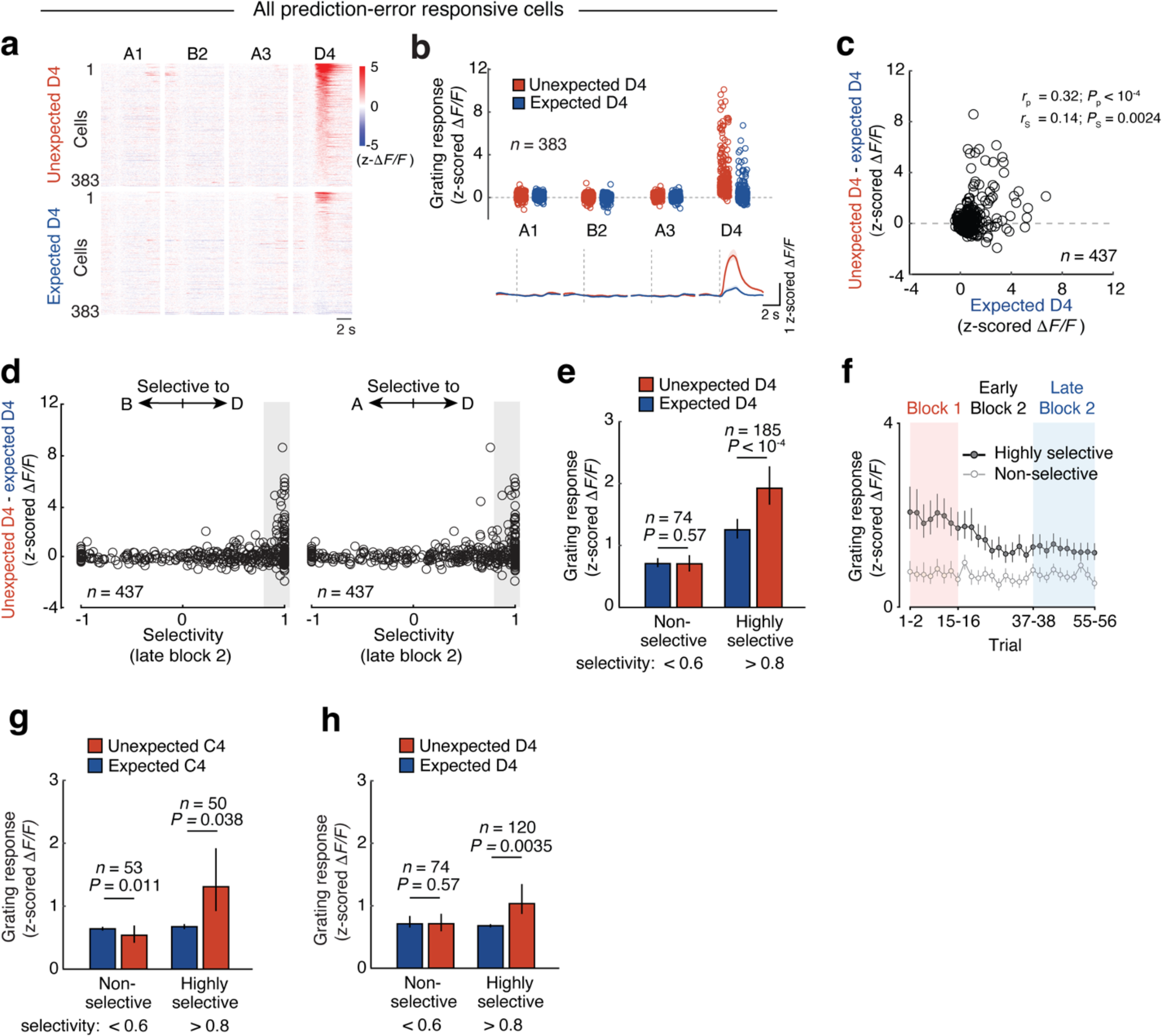
Prediction error specifically boosts neurons most selective to the presented visual stimulus (grating D). **a-f**, Same as Fig. 2, but for a second unexpected visual stimulus D. **a**, Single-cell responses for all prediction-error responsive cells (individual rows) (*n* = 383 cells, 7 mice) to gratings A1, B2, A3 and D4 in unexpected D4 (top) and expected D4 (bottom) trials, sorted by response to unexpected D4. **b**, Top: calcium responses for all prediction-error responsive cells (individual dots) (*n* = 383 cells, 7 mice) to gratings A1, B2, A3 and D4 in unexpected D4 (red) and expected D4 (blue) trials. Bottom: Cell-averaged calcium responses. Lines and shading are mean and bootstrap 95% CI. **c**, Difference in response strength between unexpected (block 1) and expected D4 (late block 2) for all grating-responsive cells in late block 2, plotted against response to expected D4 (late block 2) for individual neurons. *r*_p_ = 0.32, *r*_s_ = 0.14; *P*_p_ < 1.0 × 10^−4^, *P*_s_ = 0.0024; Pearson and Spearman correlation, respectively; *n* = 437, 7 mice. **d**, Left: difference in response strength between unexpected and expected D4 responses of individual neurons, plotted against their response selectivity to stimulus D vs. stimulus B in late block 2 (difference in response strength between expected D4 and B2 divided by the sum of responses to both stimuli) for all neurons responsive to at least one of the grating stimuli in late block 2. −1 indicates only responsive to B, +1 only responsive to D, and 0 equal responses to both. Right: same as on the left but for response selectivity to stimulus D vs. stimulus A in late block 2. **e**, Mean responses to expected (blue) and unexpected (red) stimulus D4 of non-selective (selectivity towards D, compared to B < 0.6, left, *n* = 74; *P* = 0.57; Wilcoxon signed-rank test) and highly selective (selectivity towards D, compared to B > 0.8, right, *n* = 185; *P* < 1.0 × 10^−4^) stimulus D4 responsive cells in late block 2. Error bars are 95% bootstrap CI. **f**, Mean calcium responses to stimulus D4 over all trials in the imaging session of highly selective (dark gray, *n* = 185) and non-selective (light gray, *n* = 75) stimulus D4 responsive cells in block 2. Responses were averaged over two trials. Error bars are bootstrap 95% CI. **g**, Same as Fig. 2e, but highly selective cells were sub-selected to match their average response strength to the expected stimulus C4 with the average response to expected stimulus C4 of non-selective cells. To achieve this, highly selective cells that responded strongly to expected gratings (top 35%) were removed from the analysis. **h**, Same as **g**, but for sessions with unexpected stimulus D, equivalent to panel **e**, but with matched average response strength to the expected stimulus D4 of highly selective and non-selective V1 cells.

**Extended Data Fig. 5.**
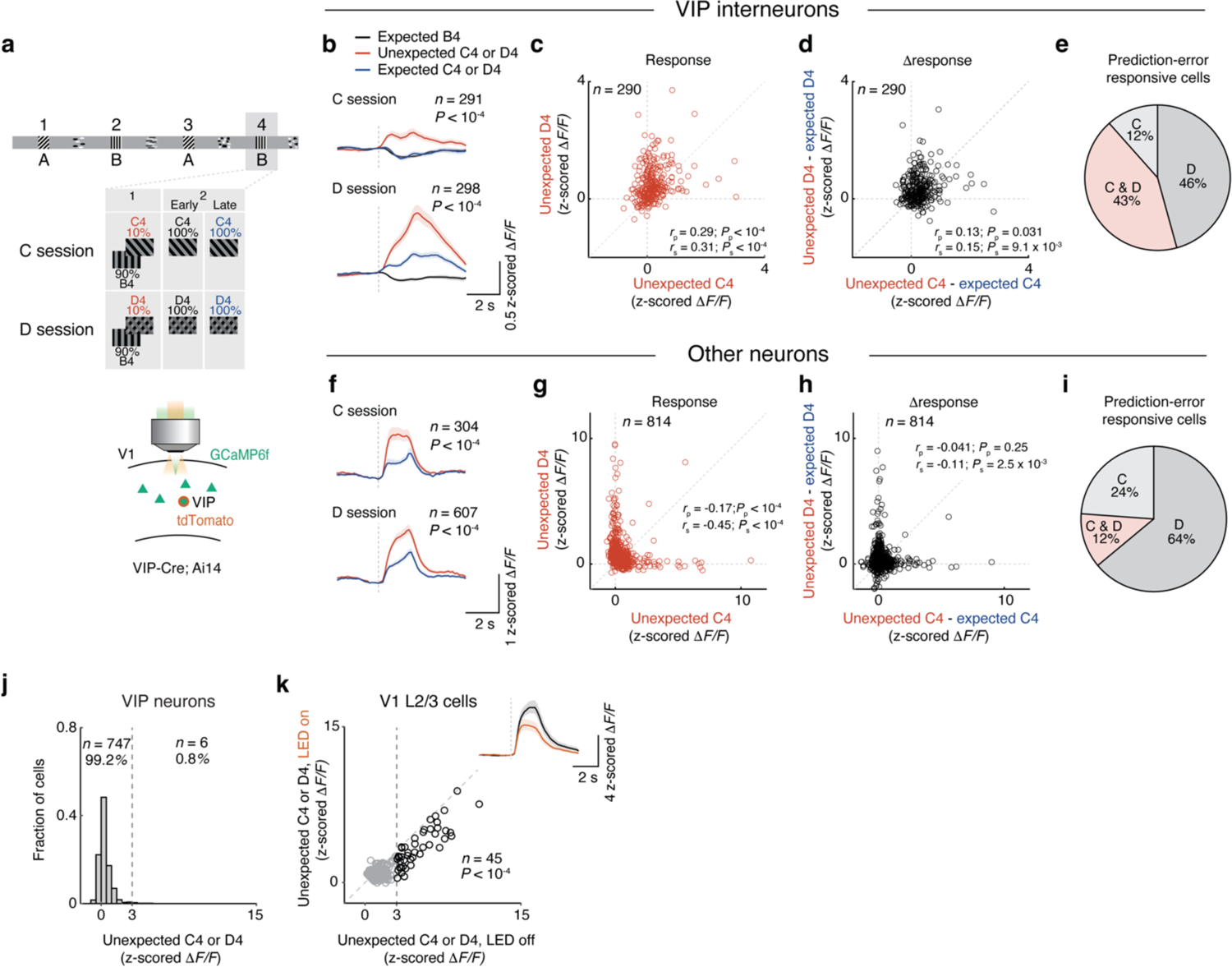
Responses of VIP and non-VIP cells. **a-i**, Same as Fig. 1k-n, but VIP neurons and other non-VIP neurons were plotted separately. **a**, Schematic of the experimental design. Stimulus C or D was presented at position 4 (C4 and D4) in 10% of trials in different sessions (C and D sessions, respectively). Calcium activity of VIP cells and other non-VIP cells in V1 layer 2/3 was recorded. **b**, Top: cell- and trial-averaged VIP calcium responses to expected B4 (black), unexpected C4 (red, block 1) and expected C4 (blue, late block 2). *n* = 291 VIP cells from 5 mice, P < 1.0 × 10^−4^, Wilcoxon signed-rank test. Bottom: cell- and trial-averaged VIP calcium responses to expected B4 (black), unexpected D4 (red) and expected D4 (blue). *n* = 298 VIP cells from 5 mice, P < 1.0 × 10^−4^, Wilcoxon signed-rank test. Lines and shading are mean and bootstrap 95% CI. **c**, Calcium responses of individual VIP neurons to unexpected D4 plotted against responses to unexpected C4. Pearson correlation: *r*_p_ = 0.29, *P*_p_ < 1.0 × 10^−4^; Spearman correlation: *r*_s_ = 0.31, *P*_s_ < 1.0 × 10^−4^, *n* = 290 from 5 mice. **d**, Difference of grating responses to unexpected and expected D4 plotted against the difference of unexpected and expected C4 (Pearson correlation: *r*_p_ = 0.13, *P*_p_ =0.031; Spearman correlation: *r*_s_ = 0.15, *P*_s_ = 9.1× 10^−3^, *n* = 290 from 5 mice. **e**, Polar plot with proportion of prediction-error responsive VIP cells for stimulus C, stimulus D, or both (see Methods). *n* = 199 from 5 mice. **f-i**, Same as **b**-**e**, but for non-VIP cells. **f**, Top: cell- and trial-averaged calcium responses of C4-responsive neurons to unexpected C4 (red, block 1) and expected C4 (blue, late block 2). *n* = 304 cells from 5 mice, *P* < 1.0 × 10^−4^, Wilcoxon signed-rank test. Bottom: cell- and trial-averaged calcium responses to unexpected D4 (red) and expected D4 (blue). *n* = 607 cells from 5 mice, *P* < 1.0 × 10^−4^, Wilcoxon signed-rank test. Lines and shading are mean and bootstrap 95% CI. **g**, Calcium responses of individual V1 neurons to unexpected D4 plotted against unexpected C4. Pearson correlation: *r*_p_ = −0.17, *P*_p_ < 1.0 × 10^−4^; Spearman correlation: *r*_s_ = −0.45, *P*_s_ < 1.0 × 10^−4^, *n* = 814 cells from 5 mice. **h**, The Difference of stimulus responses between unexpected and expected D4 plotted against the difference of responses between unexpected and expected C4. Pearson correlation: *r*_p_ = −0.041, *P*_p_ =0.25; Spearman correlation: *r*_s_ = −0.11, *P*_s_ = 2.5 × 10^−3^, *n* = 814 cells from 5 mice. **i**, Polar plot with proportion of prediction-error responsive non-VIP neurons for stimulus C, stimulus D, or both (see Methods). *n* = 960 cells from 5 mice. **j**, Distribution of stimulus response strength for VIP cells to unexpected C4 or D4 (*n* = 753, 14 sessions from 7 mice). Same data as Fig. 3a-d. **k**, Same as Fig. 3f, but only cells exhibiting a visual stimulus response of more than 3 z-scored *ΔF/F* were included in order to avoid the signal contribution of opsin-expressing and therefore directly silenced VIP cells (see panel j), which cannot be visually identified in these experiments (*n* = 45 from 7 sessions; *P* < 1.0 × 10^−4^; Wilcoxon signed-rank test). Neurons indicated in black have responses > 3 z-scored *ΔF/F.* Inset: Responses to unexpected stimulus C4 or D4 of V1 layer 2/3 cells with (amber) or without (black) VIP silencing. Lines and shading are mean and bootstrap 95% CI.

**Extended Data Fig. 6.**
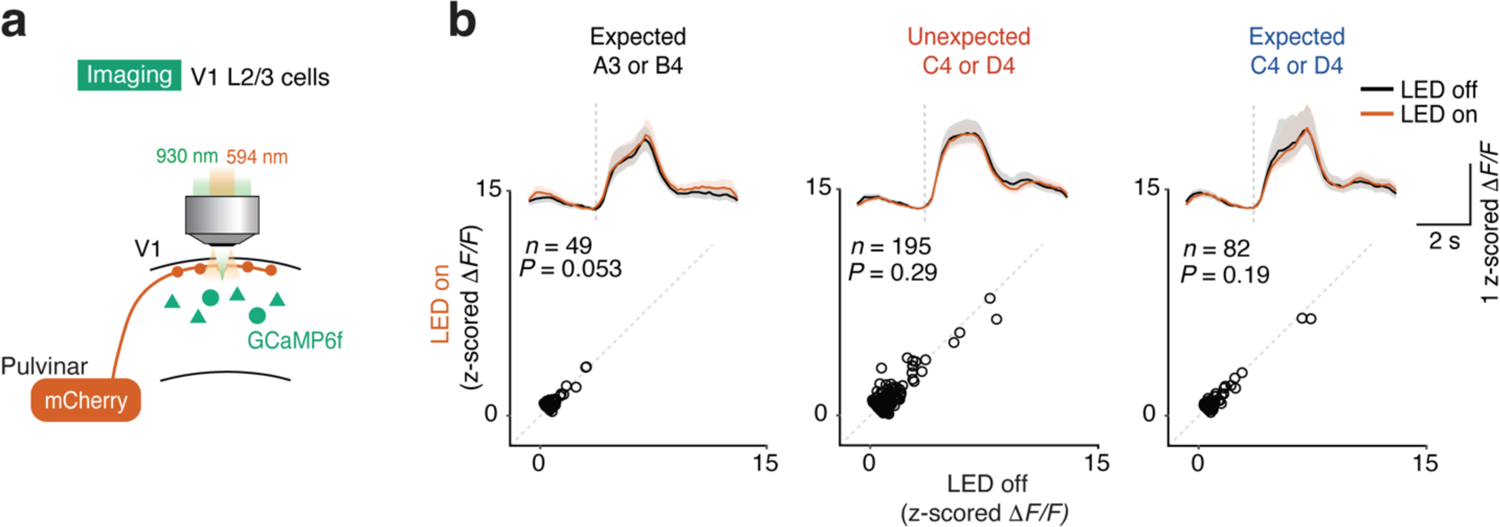
Control experiment for LED light stimulation. **a**, Schematic of the experimental design. Calcium activity of V1 layer 2/3 cells was recorded during light stimulation without expression of opsins. mCherry was expressed in pulvinar neurons. **b**, Top: cell- and trial-averaged responses to expected grating A3 or B4 (left), unexpected grating C4 or D4 (middle) and expected grating C4 or D4 (right) with or without light stimulation (amber and black, respectively). Lines and shading are mean and bootstrap 95% CI (*n* = 49, 195, 82; *P* = 0.053, *P* = 0.29, *P* = 0.19; expected grating A3 or B4 responsive cells, unexpected grating C4 or D4 responsive cells, expected grating C4 or D4 responsive cells; Wilcoxon signed-rank test; 3 mice). Bottom: Responses of individual V1 neurons to stimuli indicated above with and without LED light stimulation (LED on vs LED off).

**Extended Data Fig. 7.**
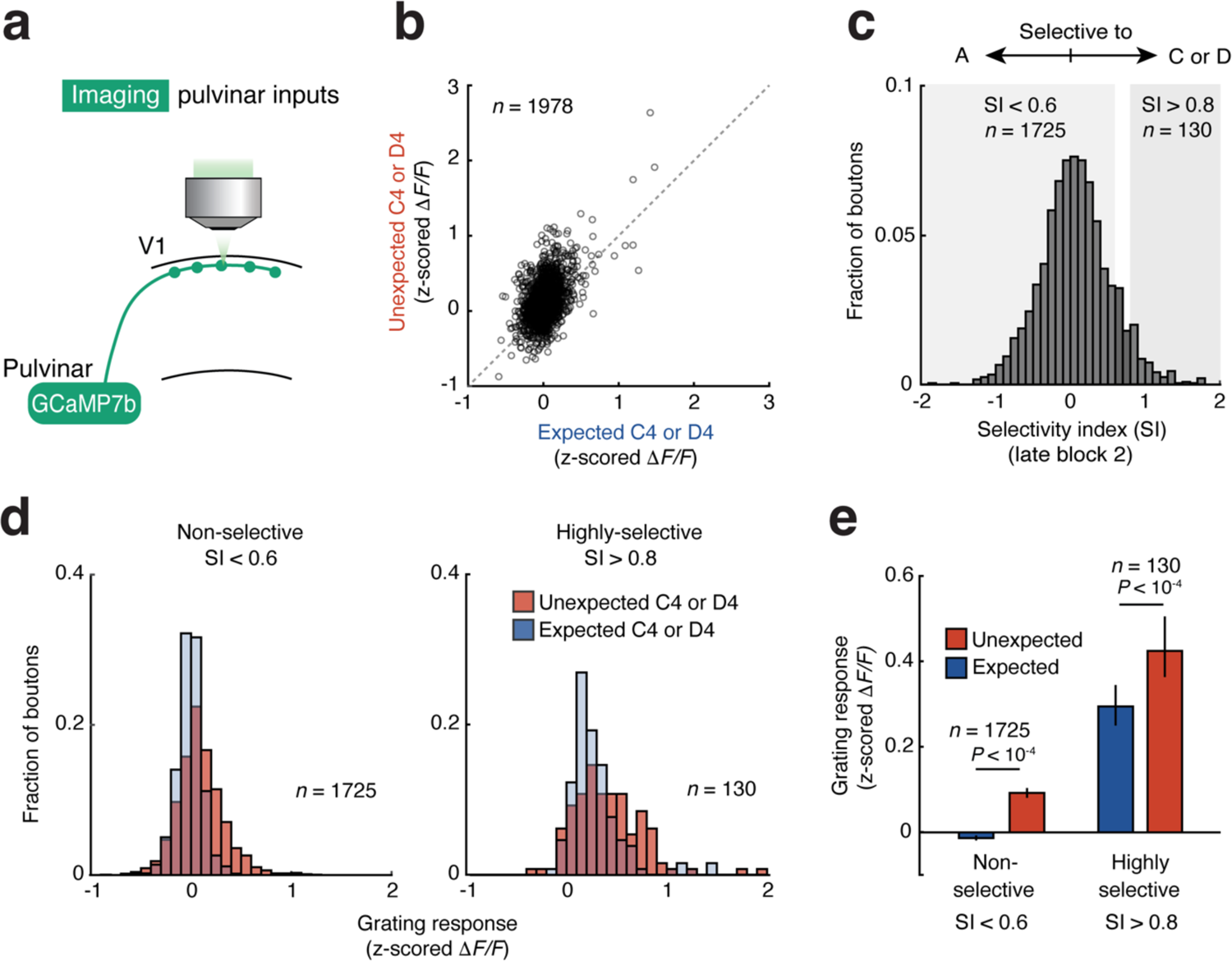
Broad facilitation of pulvinar signals by prediction errors. **a**, Experimental design. The calcium activity of axonal boutons of pulvinar projections in V1 L1 was recorded. **b**, Stimulus responses of individual pulvinar boutons to unexpected C4 or D4 plotted against responses to expected C4 or D4. *n* = 1,978 pulvinar boutons from 10 sessions, 7 mice. **c**, Distribution of selectivity index (grating C or D vs. grating A, see Methods) for all pulvinar boutons. **d**, Distribution of stimulus response strength of non-selective (selectivity index C4/D4, compared to A3 < 0.6, left) and highly selective (selectivity index > 0.8, right) pulvinar boutons to unexpected C4 or D4 (red) and expected C4 or D4 (blue). **e**, Cell- and trial-averaged stimulus responses to expected C4 or D4 (blue) and unexpected C4 or D4 (red), of non-selective (left) and highly selective (right) pulvinar boutons. *n* = 1,725 and *n* = 130; *P* < 1.0 × 10^−4^, *P* < 1.0 × 10^−4^; non-selective and highly selective cells, Wilcoxon signed-rank test. Bars and error bars are mean and 95% bootstrap CI.

**Extended Data Fig. 8.**
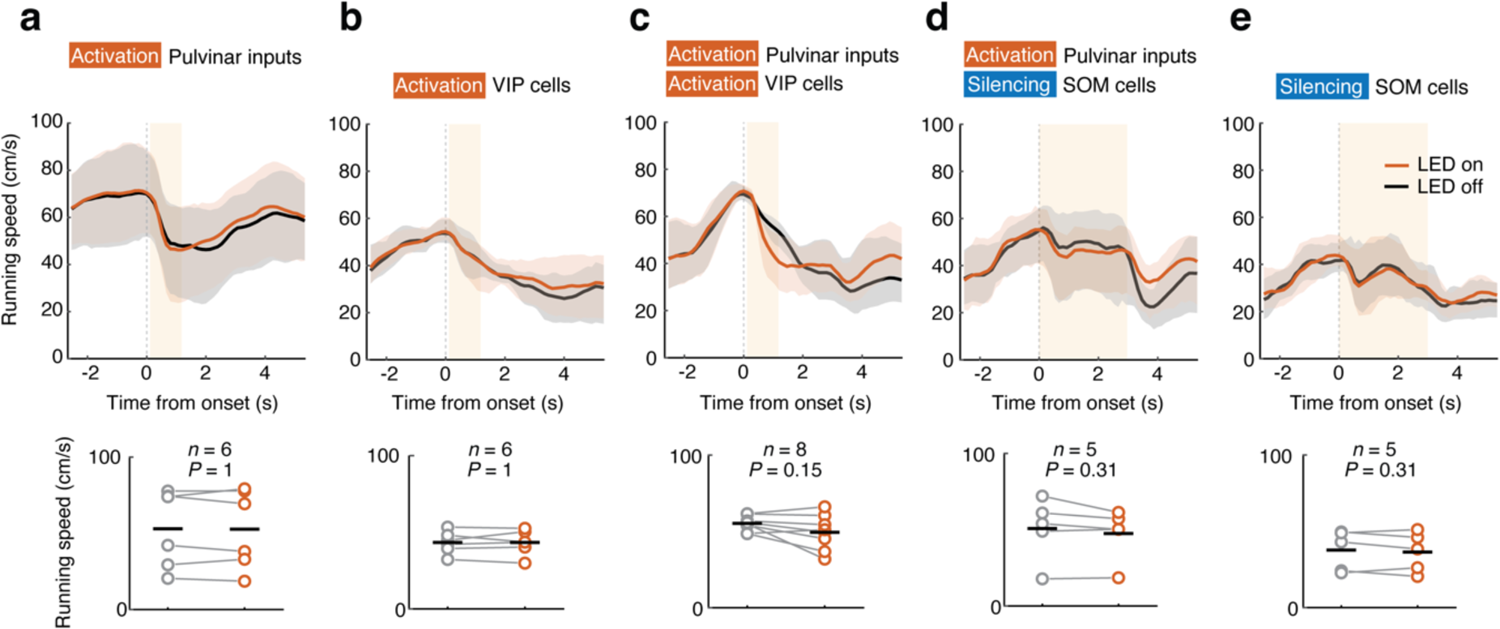
No effect on running behavior by optogenetic manipulations (related to Fig. 4 and Extended Data Fig. 10). **a-e**, Running speed with (amber) or without (black) optogenetic manipulation for activation of pulvinar axons (**a**), activation of VIP neurons (**b**), co-activation of pulvinar axons and VIP neurons (**c**), activation of pulvinar axons and simultaneous silencing of SOM cells (**d**), and silencing of SOM cells (**e**). Top: Lines and shading are mean and bootstrap 95% CI. Orange shading indicates time of optogenetic stimulation. Bottom: Data from the individual animals are shown separately. Data from the same animals are connected by lines. Black horizontal bars represent mean across animals. *P* values from Wilcoxon signed-rank test.

**Extended Data Fig. 9.**
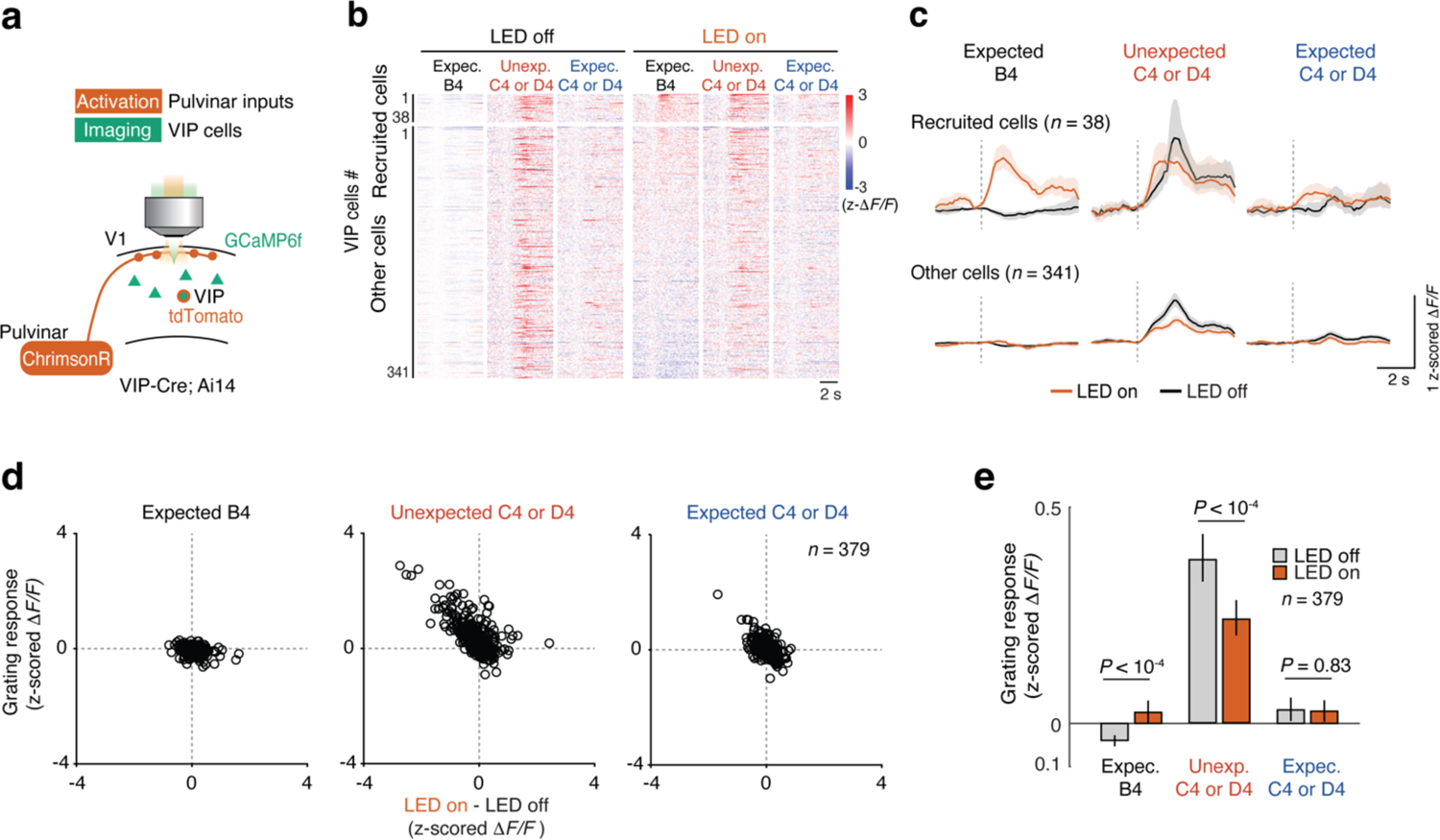
Effect of optogenetic stimulation of pulvinar inputs on grating responses of VIP cells in V1 layer 2/3. **a**, Schematic of the experimental design. The activity of VIP cells was recorded while pulvinar axons were optogenetically stimulated for 3 s, starting at the onset of the grating stimulus. **b**, Single-cell responses of pulvinar-recruited VIP cells (individual rows, *n* = 38 cells, 7 sessions from 5 mice) and other non-recruited VIP cells (individual rows, *n* = 341 cells, 7 sessions from 5 mice) to expected B4, unexpected C4 or D4 and expected C4 or D4 stimuli with (right) and without (left) optogenetic stimulation (see Methods). **c**, Cell-averaged calcium responses with (amber) or without (black) optogenetic stimulation of pulvinar-recruited and other non-recruited VIP cells. Lines and shaded regions are mean and bootstrap 95% CI. **d**, Visual stimulus responses of individual VIP neurons without optogenetic stimulation plotted against the effect of pulvinar stimulation (difference of grating responses with and without optogenetic stimulation). **e**, Responses to expected B4 (left), unexpected C4 or D4 (middle) and expected C4 or D4 (right) stimuli with (amber) and without (black) pulvinar stimulation. *n* = 379 cells from 7 sessions, 5 mice, LED on vs off during expected B4 stimulus: *P* < 1.0 × 10^-4^; LED on vs off during unexpected C4 or D4 stimulus, *P* < 1.0 × 10^-4^; LED on vs off during expected C4 or D4 stimulus, *P* = 0.83; Wilcoxon signed-rank test. Bars and error bars are mean and 95% bootstrap CI.

**Extended Data Fig. 10.**
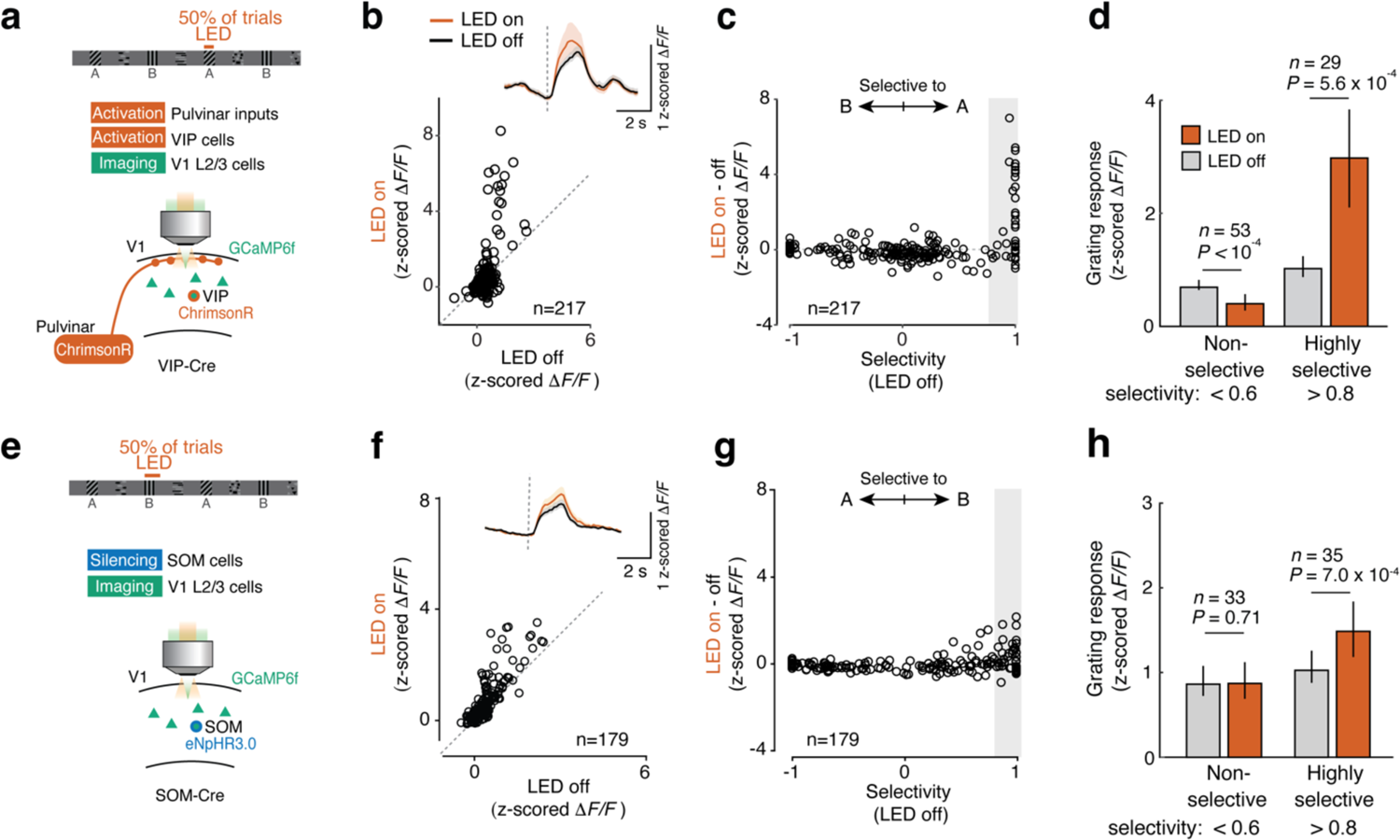
Effect of optogenetic manipulation of pulvinar inputs, VIP cells and SOM cells on grating responses of V1 layer 2/3 cells (related to Fig. 4 and Extended Data Fig. 8). **a-d**, Same as Fig. 4d, but optogenetic stimulation was paired with the grating stimulus A3 instead of B2. **a**, Schematic of the experimental design. The activity of V1 layer 2/3 cells was recorded while pulvinar axons and VIP cells were optogenetically co-stimulated. Stimulation started 0.1 s after grating onset and lasted for 1 s (see methods). **b**, Response strength to grating stimulus A3 with and without co-stimulation of pulvinar inputs and VIP cells. *n* = 217 grating A or B responsive cells, 6 sessions from 6 mice, *P* = 0.034, Wilcoxon signed-rank test. Inset: Cell-averaged calcium responses with (amber) or without (black) optogenetic stimulation. **c**, Effect of optogenetic stimulation (difference of response to grating A3 with and without laser stimulation) plotted against response selectivity (difference in response strength between stimulus A and B divided by the sum of responses) of individual V1 neurons. **d**, Calcium response strength to grating stimulus A3 of non-selective (selectivity A vs B < 0.6, left, *P* < 1.0 × 10^-4^; *n* = 53) and highly selective (selectivity B vs A > 0.8, right, *P* = 5.6 × 10^-4^; n = 29) grating A3 responsive cells in V1 layer 2/3 with (amber) or without (gray) optogenetic stimulation. Bars and error bars are mean and bootstrap 95% CI. **e-h**, Same as **a-d**, but for optogenetic silencing of SOM cells during presentation of grating stimulus B2. **e**, Schematic of the experimental design. The activity of V1 layer 2/3 cells was recorded while SOM cells were ontogenetically silenced for 3 s, starting at grating stimulus onset. **f**, Grating B2 responses with and without the silencing of SOM cells. *n* = 179 grating A or B responsive cells, 5 sessions from 3 mice, *P* = 4.0 × 10^-4^, Wilcoxon signed-rank test. **g**, Effect of optogenetic stimulation (difference of response to grating B2 with and without laser stimulation) plotted against response selectivity (difference in response strength between stimulus B and A divided by the sum of responses) of individual V1 neurons. **h**, Grating B2 responses of non-selective (selectivity B vs A < 0.6, left, *P* = 0.71; *n* = 33, Wilcoxon signed-rank test) and highly selective (selectivity B vs A > 0.8, right, *P* = 7.0 × 10^-4^; n = 35) grating B2 responsive cells in V1 layer 2/3 with (amber) or without (gray) optogenetic stimulation. Bars and error bars are mean and bootstrap 95% CI.

## References

1. Schultz, W. & Dickinson, A. Neuronal coding of prediction errors. Annu. Rev. Neurosci. 23, 473–500 (2000).

2. Starkweather, C. K., Babayan, B. M., Uchida, N. & Gershman, S. J. Dopamine reward prediction errors reflect hidden-state inference across time. Nat. Neurosci. 20, 581–589 (2017).

3. Lowet, A. S., Zheng, Q., Matias, S., Drugowitsch, J. & Uchida, N. Distributional Reinforcement Learning in the Brain. Trends Neurosci. 43, 980–997 (2020).

4. Wolpert, D. M., Miall, R. C. & Kawato, M. Internal models in the cerebellum. Trends Cogn. Sci. 2, 338–347 (1998).

5. Mumford, D. On the computational architecture of the neocortex. II. The role of cortico-cortical loops. Biol. Cybern. 66, 241–251 (1992).

6. Rao, R. P. & Ballard, D. H. Predictive coding in the visual cortex: a functional interpretation of some extra-classical receptive-field effects. Nat. Neurosci. 2, 79–87 (1999).

7. Friston, K. A theory of cortical responses. Philos. Trans. R. Soc. Lond. B Biol. Sci. 360, 815–836 (2005).

8. Clark, A. Whatever next? Predictive brains, situated agents, and the future of cognitive science. Behav. Brain Sci. 36, 181–204 (2013).

9. den Ouden, H. E. M., Kok, P. & de Lange, F. P. How prediction errors shape perception, attention, and motivation. Front. Psychol. 3, 548 (2012).

10. Keller, G. B. & Mrsic-Flogel, T. D. Predictive Processing: A Canonical Cortical Computation. Neuron 100, 424–435 (2018).

11. Rust, N. C. & Cohen, M. R. Priority coding in the visual system. Nat. Rev. Neurosci. 23, 376–388 (2022).

12. Alink, A., Schwiedrzik, C. M., Kohler, A., Singer, W. & Muckli, L. Stimulus predictability reduces responses in primary visual cortex. J. Neurosci. 30, 2960–2966 (2010).

13. Meyer, T. & Olson, C. R. Statistical learning of visual transitions in monkey inferotemporal cortex. Proc. Natl. Acad. Sci. U. S. A. 108, 19401–19406 (2011).

14. Fiser, A. et al. Experience-dependent spatial expectations in mouse visual cortex. Nat. Neurosci. 19, 1658–1664 (2016).

15. Attinger, A., Wang, B. & Keller, G. B. Visuomotor Coupling Shapes the Functional Development of Mouse Visual Cortex. Cell 169, 1291–1302.e14 (2017).

16. Audette, N. J., Zhou, W., La Chioma, A. & Schneider, D. M. Precise movement-based predictions in the mouse auditory cortex. Curr. Biol. 32, 4925–4940.e6 (2022).

17. Kim, H. R. et al. A Unified Framework for Dopamine Signals across Timescales. Cell 183, 1600–1616.e25 (2020).

18. Wessel, J. R. & Aron, A. R. On the Globality of Motor Suppression: Unexpected Events and Their Influence on Behavior and Cognition. Neuron 93, 259–280 (2017).

19. Chen, T.-W. et al. Ultrasensitive fluorescent proteins for imaging neuronal activity. Nature 499, 295–300 (2013).

20. Ranganath, C. & Rainer, G. Neural mechanisms for detecting and remembering novel events. Nat. Rev. Neurosci. 4, 193–202 (2003).

21. Marina Garrett*, Sahar Manavi, Kate Roll, Douglas R Ollerenshaw, Peter A Groblewski, Nicholas D Ponvert, Justin T Kiggins, Linzy Casal, Kyla Mace, Ali Williford, Arielle Leon, Xiaoxuan Jia, Peter Ledochowitsch, Michael A Buice, Wayne Wakeman, Stefan Mihalas, Shawn R Olsen*. Experience shapes activity dynamics and stimulus coding of VIP inhibitory cells. Elife.

22. Garrett, M., et al. Stimulus novelty uncovers coding diversity in visual cortical circuits. *bioRxiv* 2023.02.14.528085 (2023) doi:10.1101/2023.02.14.528085.

23. Homann, J., Koay, S. A., Chen, K. S., Tank, D. W. & Berry, M. J. Novel stimuli evoke excess activity in the mouse primary visual cortex. Proceedings of the National Academy of Sciences 119, e2108882119 (2022).

24. Tang, M. F. et al. Expectation violations enhance neuronal encoding of sensory information in mouse primary visual cortex. Nat. Commun. 14, 1196 (2023).

25. Pi, H.-J. et al. Cortical interneurons that specialize in disinhibitory control. Nature 503, 521–524 (2013).

26. Pfeffer, C. K., Xue, M., He, M., Huang, Z. J. & Scanziani, M. Inhibition of inhibition in visual cortex: the logic of connections between molecularly distinct interneurons. Nat. Neurosci. 16, 1068–1076 (2013).

27. Lee, S., Kruglikov, I., Huang, Z. J., Fishell, G. & Rudy, B. A disinhibitory circuit mediates motor integration in the somatosensory cortex. Nat. Neurosci. 16, 1662–1670 (2013).

28. Fu, Y. et al. A cortical circuit for gain control by behavioral state. Cell 156, 1139–1152 (2014).

29. Schneider-Mizell, C. M. et al. Cell-type-specific inhibitory circuitry from a connectomic census of mouse visual cortex. bioRxiv (2023) doi:10.1101/2023.01.23.525290.

30. Zhang, S. et al. Long-range and local circuits for top-down modulation of visual cortex processing. Science 345, 660–665 (2014).

31. Ma, G. et al. Hierarchy in sensory processing reflected by innervation balance on cortical interneurons. Sci Adv 7, (2021).

32. Roth, M. M. et al. Thalamic nuclei convey diverse contextual information to layer 1 of visual cortex. Nat. Neurosci. 19, 299–307 (2016).

33. Blot, A. et al. Visual intracortical and transthalamic pathways carry distinct information to cortical areas. Neuron 109, 1996–2008.e6 (2021).

34. Bennett, C. et al. Higher-Order Thalamic Circuits Channel Parallel Streams of Visual Information in Mice. Neuron 102, 477–492.e5 (2019).

35. Harris, J. A. et al. Hierarchical organization of cortical and thalamic connectivity. Nature 575, 195–202 (2019).

36. Petty, G. H., Kinnischtzke, A. K., Hong, Y. K. & Bruno, R. M. Effects of arousal and movement on secondary somatosensory and visual thalamus. Elife 10, (2021).

37. Fang, Q. et al. A Differential Circuit via Retino-Colliculo-Pulvinar Pathway Enhances Feature Selectivity in Visual Cortex through Surround Suppression. Neuron 105, 355–369.e6 (2020).

38. Audette, N. J., Urban-Ciecko, J., Matsushita, M. & Barth, A. L. POm Thalamocortical Input Drives Layer-Specific Microcircuits in Somatosensory Cortex. Cereb. Cortex 28, 1312–1328 (2018).

39. Sermet, B. S. et al. Pathway-, layer- and cell-type-specific thalamic input to mouse barrel cortex. Elife 8, (2019).

40. Adesnik, H., Bruns, W., Taniguchi, H., Huang, Z. J. & Scanziani, M. A neural circuit for spatial summation in visual cortex. Nature 490, 226–231 (2012).

41. Pala, A. & Petersen, C. C. H. In vivo measurement of cell-type-specific synaptic connectivity and synaptic transmission in layer 2/3 mouse barrel cortex. Neuron 85, 68–75 (2015).

42. Kanai, R., Komura, Y., Shipp, S. & Friston, K. Cerebral hierarchies: predictive processing, precision and the pulvinar. Philos. Trans. R. Soc. Lond. B Biol. Sci. 370, (2015).

43. Hu, F. et al. Prefrontal Corticotectal Neurons Enhance Visual Processing through the Superior Colliculus and Pulvinar Thalamus. Neuron 104, 1141–1152.e4 (2019).

44. Melzer, S. et al. Bombesin-like peptide recruits disinhibitory cortical circuits and enhances fear memories. Cell 184, 5622–5634.e25 (2021).

45. Szadai, Z. et al. Cortex-wide response mode of VIP-expressing inhibitory neurons by reward and punishment. Elife 11, (2022).

46. Ren, C. et al. Global and subtype-specific modulation of cortical inhibitory neurons regulated by acetylcholine during motor learning. Neuron 110, 2334–2350.e8 (2022).

47. Cossell, L. et al. Functional organization of excitatory synaptic strength in primary visual cortex. Nature 518, 399–403 (2015).

48. Znamenskiy, P., Kim, M. H., Muir, D. R. & Iacaruso, M. F. Functional selectivity and specific connectivity of inhibitory neurons in primary visual cortex. Biorxiv (2018).

49. Bock, D. D. et al. Network anatomy and in vivo physiology of visual cortical neurons. Nature 471, 177–182 (2011).

50. Packer, A. M. & Yuste, R. Dense, unspecific connectivity of neocortical parvalbumin-positive interneurons: a canonical microcircuit for inhibition? J. Neurosci. 31, 13260– 13271 (2011).

51. Hangya, B., Ranade, S. P., Lorenc, M. & Kepecs, A. Central Cholinergic Neurons Are Rapidly Recruited by Reinforcement Feedback. Cell 162, 1155–1168 (2015).

52. Jordan, R. & Keller, G. B. The locus coeruleus broadcasts prediction errors across the cortex to promote sensorimotor plasticity. *bioRxiv* 2022.11.08.515698 (2023) doi:10.1101/2022.11.08.515698.

53. Pologruto, T. A., Sabatini, B. L. & Svoboda, K. ScanImage: flexible software for operating laser scanning microscopes. Biomed. Eng. Online 2, 13 (2003).

54. Leinweber, M. et al. Two-photon calcium imaging in mice navigating a virtual reality environment. J. Vis. Exp. e50885 (2014) doi:10.3791/50885.

55. Muir, D. R., Roth, M. & Blot, A. TimeSeries analysis toolbox for Matlab. (2020) doi:10.5281/zenodo.3859433.

56. ast_model: Asymmetric Student-t model for neuropil decontamination. (Github).

57. Poort, J. et al. Learning Enhances Sensory and Multiple Non-sensory Representations in Primary Visual Cortex. Neuron 86, 1478–1490 (2015).

